# An Empirical Bayes Approach to Estimating Dynamic Models of Co-Regulated Gene Expression

**DOI:** 10.1101/2021.07.08.451684

**Authors:** Sara Venkatraman, Sumanta Basu, Andrew G. Clark, Sofie Delbare, Myung Hee Lee, Martin T. Wells

## Abstract

Time-course gene expression datasets provide insight into the dynamics of complex biological processes, such as immune response and organ development. It is of interest to identify genes with similar temporal expression patterns because such genes are often biologically related. However, this task is challenging due to the high dimensionality of these datasets and the nonlinearity of gene expression time dynamics. We propose an empirical Bayes approach to estimating ordinary differential equation (ODE) models of gene expression, from which we derive a similarity metric between genes called the Bayesian lead-lag *R*^2^ (LL*R*^2^). Importantly, the calculation of the LL*R*^2^ leverages biological databases that document known interactions amongst genes; this information is automatically used to define informative prior distributions on the ODE model’s parameters. As a result, the LL*R*^2^ is a biologically-informed metric that can be used to identify clusters or networks of functionally-related genes with co-moving or time-delayed expression patterns. We then derive data-driven shrinkage parameters from Stein’s unbiased risk estimate that optimally balance the ODE model’s fit to both data and external biological information. Using real gene expression data, we demonstrate that our methodology allows us to recover interpretable gene clusters and sparse networks. These results reveal new insights about the dynamics of biological systems.

## 1 Introduction

Time-course gene expression datasets are an essential resource for querying the dynamics of complex biological processes, such as immune response, disease progression, and organ development [Yosef and Regev, 2011, Bar-Joseph et al., 2012, Purvis and Lahav, 2013]. Such datasets, now abundantly available through techniques such as whole-genome RNA sequencing, consist of gene expression measurements for thousands of an organism’s genes at a few (typically 5-20) time points. Experimental evidence has revealed that groups of genes exhibiting similar temporal expression patterns are often biologically associated [Eisen et al., 1998]. For instance, such genes may be co-regulated by the same *transcription factors* [Tavazoie et al., 1999]: proteins that directly control gene expression, which ultimately contributes to changes in cellular function. Identifying clusters or networks of genes with related temporal dynamics, which is our objective in this study, can therefore uncover the regulators of dynamic biological processes. Doing so can also help generate testable hypotheses about the roles of orphan genes that exhibit similar expression patterns to ones that are better understood.

The complex, nonlinear time dynamics of gene expression pose a significant challenge for clustering and network analysis in genomics. Groups of interacting genes may be expressed with time lags or inverted patterns [Qian et al., 2001] due to delayed activation of underlying transcription factors, making it difficult to measure the “similarity” in two expression profiles. Ordinary differential equations (ODEs) or discrete-time difference equations have been successfully used to model the nonlinear expressions of a small number of genes [D’haeseleer et al., 1999, Chen et al., 1999, De Jong, 2002, Bansal et al., 2006, Polynikis et al., 2009]. It is possible to derive similarity metrics for the time dynamics of two genes from such ODEs, thus enabling putative identification of co-regulated genes and the reconstruction of regulatory networks [Farina et al., 2007, 2008, Wu et al., 2019]. In particular, the approach proposed in Farina et al. [2007] allows explicit modeling of lead-lag as well as contemporaneous associations between gene expression trajectories. We hence use it as the basis of the similarity calculations in our proposed clustering framework.

The high dimensionality (number of genes) and small sample sizes (number of time points) of time-course gene expression datasets pose another obstacle to identifying genes with similar expression dynamics. Due to the size of these datasets, the number of gene pairs receiving high similarity scores by any method can be overwhelmingly large. High similarity scores are typically validated for biological relevance using annotations provided by extensive public and commercial curated databases that assign genes to functional groups. For instance, Gene Ontology (GO) annotations are keywords that describe a gene’s molecular function, role in a biological process, or cellular localization [Ashburner et al., 2000]. Other curated databases include KEGG [Kanehisa and Goto, 2000], Reactome [Fabregat et al., 2018], BioCyc [Karp et al., 2019], and STRING [Szklarczyk et al., 2019]. To ease the burden of manually validating a potentially vast number of gene-gene associations, we propose a Bayesian clustering technique that uses annotations as prior information to automatically validate these associations. Incorporating such information into a clustering method can encourage gene pairs with known biological associations to receive higher similarity scores, while filtering away those known to be unrelated. This also allows for knowledge gleaned from gene expression time series data to be contrasted with other knowledge bases; for instance, two genes with highly similar temporal expression patterns may not have been considered associated in previous cross-sectional (single time point) studies on which annotations are based, or vice versa.

There exist in the literature a few approaches to integrating biological knowledge with statistical measures of genetic association. One line of research considers Bayesian methods that use external data sources to determine prior distributions over genes or proteins that influence a biological response [Li and Zhang, 2010, Stingo et al., 2011, Hill et al., 2012, Lo et al., 2012, Peng et al., 2013]. Other studies develop biologically-informed regularization terms in graph-regularized methods for reconstructing gene networks [Zhang et al., 2013, Li and Jackson, 2015]. In another work, Nepomuceno et al. [2015] propose an algorithm for biclustering gene expression data using gene ontology annotations. However, less attention has been given to using both data and prior biological knowledge to identify and model dominant patterns in the complex temporal dynamics of gene expression, e.g. with ODEs.

Our technical contribution in this work is a Bayesian method for constructing biologically-meaningful clusters and networks of genes from time-course expression data, using a new similarity measure between two genes called the *Bayesian lead-lag R*^2^ (LL*R*^2^). The Bayesian LL*R*^2^ is derived from ODE models of temporal gene expression, and is based on associations in both the time-course data and prior biological annotations. The balance between data and prior information is controlled by data-driven hyperparameters, making our approach an empirical Bayes method. As indicated by the name, the Bayesian LL*R*^2^ is based on the familiar *R*^2^ statistic (the coefficient of determination) and is simple and fast to compute for all 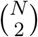 gene pairs, where *N* is the number of genes under study. Importantly, external biological information regularizes the set of significant gene-gene associations found within a time-course dataset. In Figure 1, for instance, we present a network of 1735 genes in *Drosophila melanogaster* (fruit fly) constructed both without external information, using an ordinary least-squares version of the LL*R*^2^ proposed by Farina et al. [2008], and with external information, using our proposed Bayesian LL*R*^2^; the latter is a noticeably sparsified revision of the former, and retains only edges connecting genes with either known or highly plausible biological relationships.

**Figure 1:**
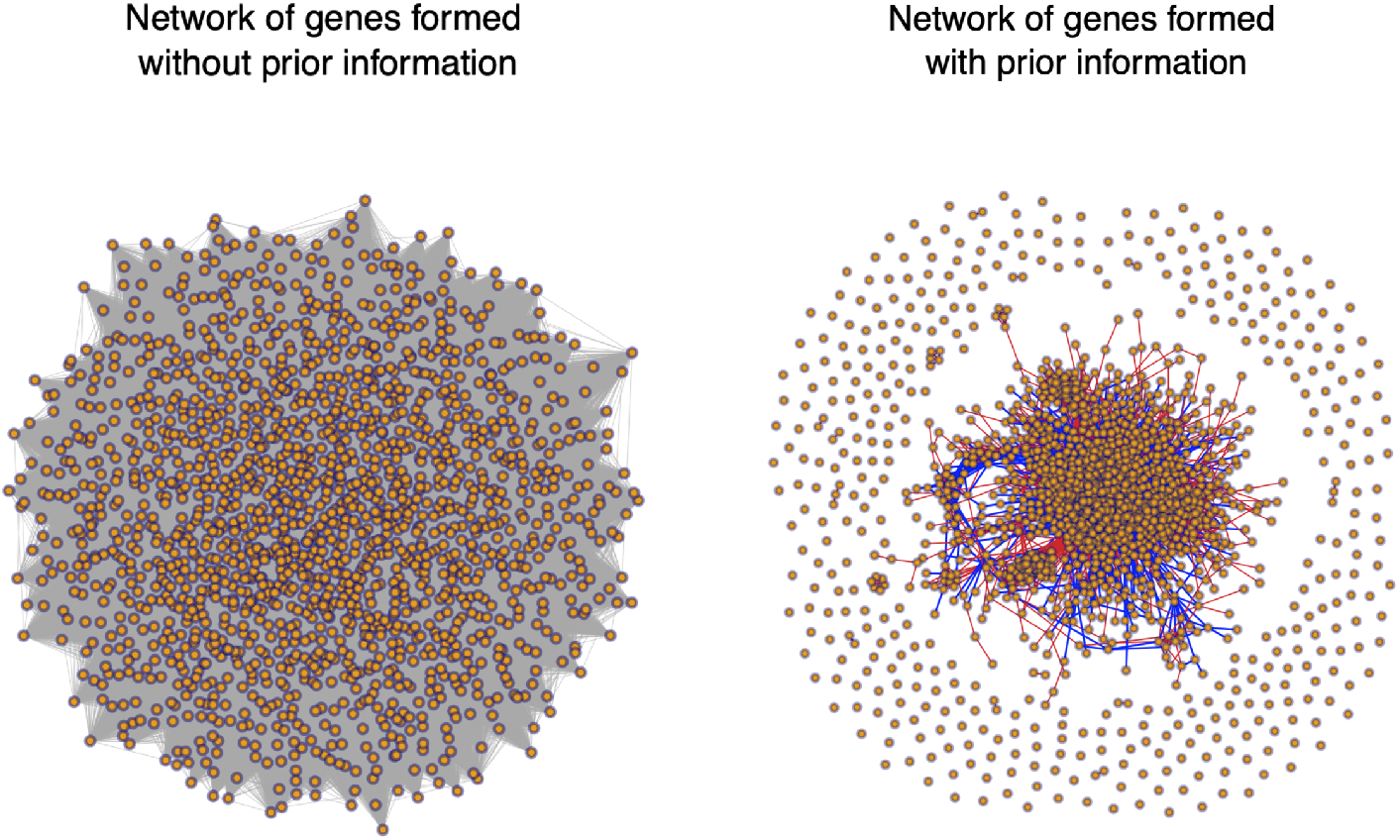
Networks of 1735 genes profiled in a time-course gene expression dataset collected by Schlamp et al. [2021]. Vertices represent genes and edges connect two genes if their lead-lag R^2^ exceeds 0.9. Left: lead-lag R^2^ is computed using ordinary least squares regression according to Farina et al. [2008], without any external biological information. All 1735 genes form a single connected component (599,896 edges). Right: lead-lag R^2^ is computed using our proposed Bayesian approach, which leverages external sources of biological information about gene-gene relationships. Red edges (11,380 edges) connect genes known to be associated. Blue edges (2830 edges) connect genes whose relationship is unknown but is supported by the data.

The remainder of this paper is organized as follows. Section 2 describes the ODE model of temporal gene expression that we adopt. Section 3 details our empirical Bayes method for fitting the ODE model and obtaining our proposed LL*R*^2^ metric. Section 4 demonstrates the application of our method to real gene expression data collected by Schlamp et al. [2021]. We recover a tradeoff between immune response and metabolism that has been observed in several studies and present examples of biologically-meaningful gene clusters identified with the Bayesian LL*R*^2^. We also discuss the method’s potential to conjecture new data-driven hypotheses about gene-gene interactions. Sections 4.2 and Appendices D-E compare the Bayesian LL*R*^2^ to its non-Bayesian counterpart as well as the commonly used Pearson correlation between genes. Proofs and additional examples are provided in the Appendix.

## 2 Dynamic models of gene expression

### 2.1 Model derivation

We consider an ODE model of gene expression proposed by Farina et al. [2007]. Let *m*_*A*_(*t*) denote the expression of some gene *A* at time *t*, measured for instance as the log_2_-fold change in mRNA levels relative to time 0. The model assumes the rate of change in gene *A*’s expression is given by some regulatory signal *p*(*t*):

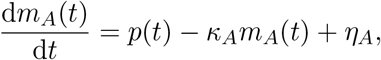

where *κ*_*A*_ denotes the mRNA decay rate for gene *A*, and *η*_*A*_ denotes natural and experimental noise. This model can be naturally extended to consider two genes *A* and *B* that might be associated with one another, i.e. that are governed by the same underlying *p*(*t*), yielding a pair of coupled differential equations:

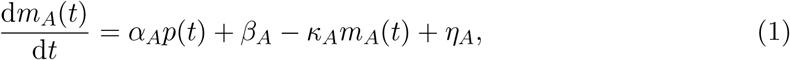

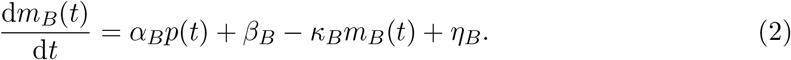

The common signal *p*(*t*) accounts for the effect of one or more transcription factors that potentially regulate both genes *A* and *B*. The coefficients *α*_*A*_ and *α*_*B*_ measure the strength of *p*(*t*) in the expression patterns of genes *A* and *B*, respectively. *β*_*A*_ and *β*_*B*_ are affine coefficients allowing *m*_*A*_(*t*) and *m*_*B*_(*t*) to exhibit linear time trends.

We now obtain a model of gene *A*’s expression in terms of gene *B*’s expression by first rearranging (2) to isolate *p*(*t*):

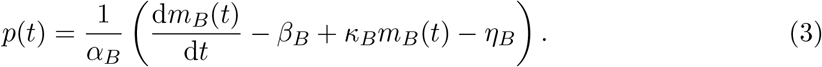

Substituting (3) into (1) yields

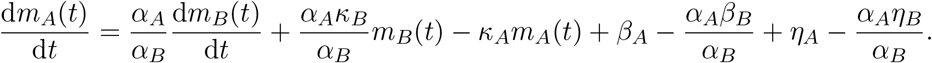

Integrating from 0 to *t*, we obtain:

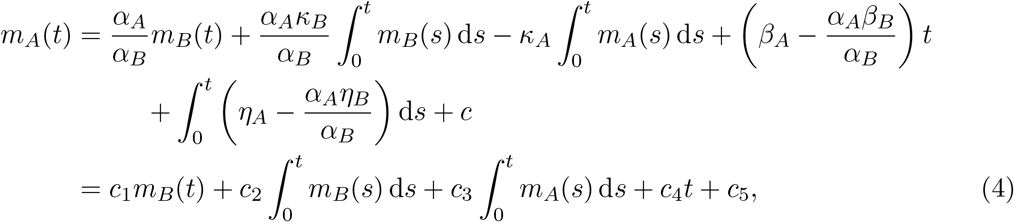

where 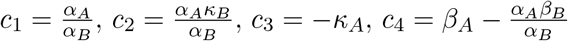, and 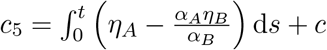.

Observe that (4) is linear in the parameters *c*_*k*_. Thus, given measurements {*m*_*A*_(*t*_1_), …, *m*_*A*_(*t*_*n*_)} and {*m*_*B*_(*t*_1_), …, *m*_*B*_(*t*_*n*_)} of the expression levels of genes *A* and *B* at time points *t*_1_, …, *t*_*n*_,

we can express (4) as the linear model **Y** = **X*β*** + ***ε***, where

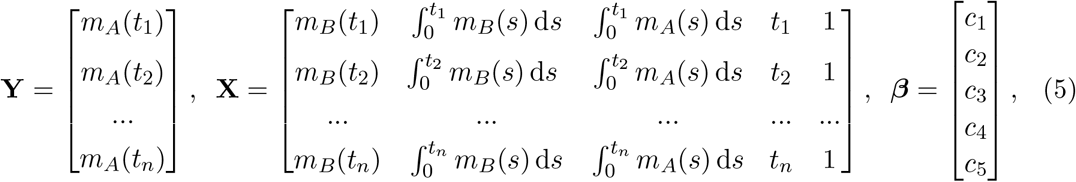

with the standard assumption that ***ε*** ∼ *N* (**0**, *σ*^2^**I**_*n*_), where **I**_*n*_ denotes the *n*×*n* identity matrix. Although only samples from the functions *m*_*A*_(*t*) and *m*_*B*_(*t*) are given, we can estimate the integral entries of the second and third columns of **X** by numerically integrating spline or polynomial interpolants fitted to these samples.

In fitting the model (4) to the expression data of genes *A* and *B*, we obtain the ordinary least-squares (OLS) estimator 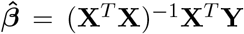. The amount of variation in gene *A*’s expression that is captured by the estimated linear model is expressed as the familiar *R*^2^ value: 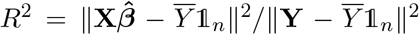, where 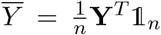 and 𝕝_*n*_ denotes the *n* × 1 vector of ones. Adopting the terminology in Farina et al. [2008], we refer to this *R*^2^ as the *lead-lag R*^2^ *between genes A and B*. A high lead-lag *R*^2^ may indicate that the two genes are co-regulated in time by common transcription factors, or are at least associated with one another in some way. The term “lead-lag” comes from the lead-lag compensator in control theory. In this context, a “lead-lag relationship” between genes refers to the presence of a common regulatory signal (input) that, in conjunction with the process of mRNA decay, modulates the expression of genes with the same biological function (output).

### 2.2 Motivating the Bayesian lead-lag *R*^2^

Our primary contribution in this work is a biologically-informed method for clustering genes based on their temporal dynamics. Clustering involves measuring the similarity between two objects, which can also be thought of as defining an edge between two nodes in an undirected network. Our similarity measure is a Bayesian version of the lead-lag *R*^2^ that uses both temporal expression data for genes *A* and *B* as well as a prior indication of whether they are biologically associated.

We can motivate the Bayesian lead-lag *R*^2^ via Figure 2, which shows examples of *false positive* gene pairs: genes that have a spuriously high lead-lag *R*^2^, but do not have similar expression patterns nor a biological relationship. The data comes from an experiment on fruit flies by Schlamp et al. [2021] that aimed to profile the dynamics of genes involved in or affected by immune response immediately following an infection. More details on this dataset can be found in Section 4.1.

**Figure 2:**
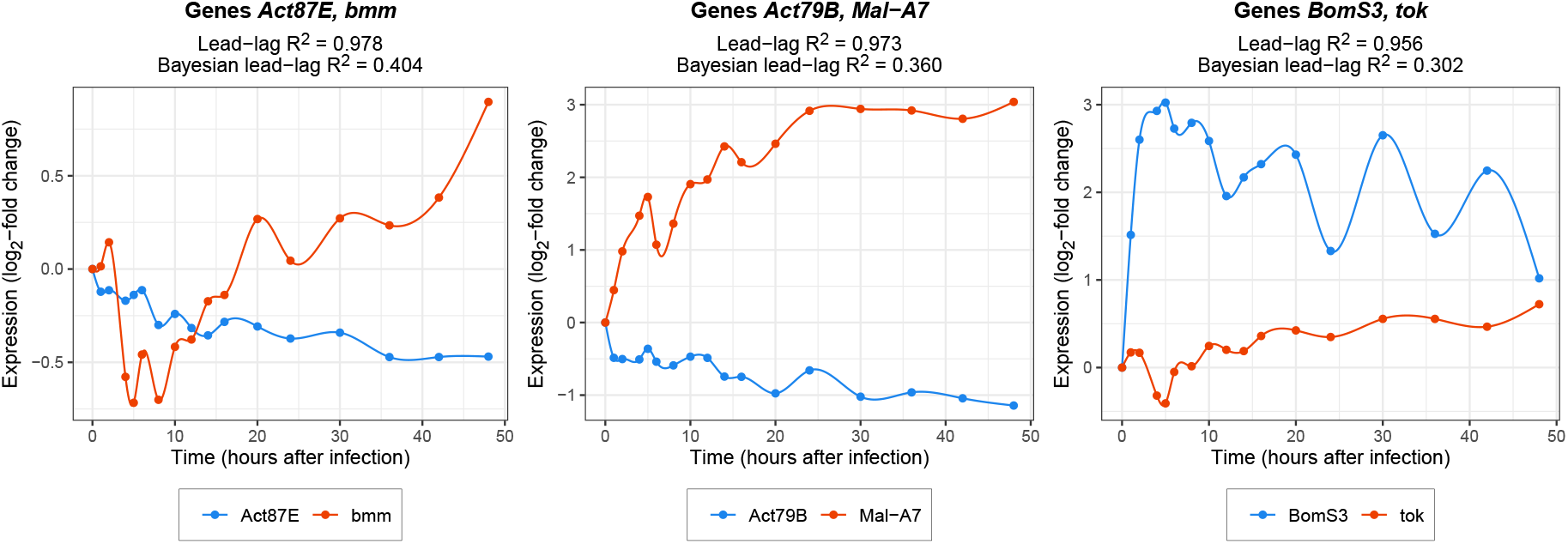
Plots of the temporal expression profiles of three gene pairs for which the lead-lag R^2^ is spuriously high. Spline interpolants were fit through the observed points, which are indicated by solid dots. Functional annotations for these genes in Flybase [Larkin et al., 2020] do not suggest a clear link within each pair. By contrast, the lower Bayesian lead-lag R^2^ values more accurately reflect the degree of these associations.

Spuriously high lead-lag *R*^2^ values are likely to arise in large datasets. For example, if gene *A*’s expression levels increase or decrease monotonically with time, the response vector **Y** in (5) will be highly correlated with the time integrals and time points in the third and fourth columns of **X**. The lead-lag *R*^2^ between genes *A* and *B* will be large, but not because the genes are associated either in time or biologically.

In contrast to gene pairs of the kind shown in Figure 2, we can consider Figure 3, which displays genes from two well-studied functional groups, known as *pathways*: circadian rhythms and immune response. Within each group, we expect to see high pairwise lead-lag *R*^2^ values (*true positives*).

**Figure 3:**
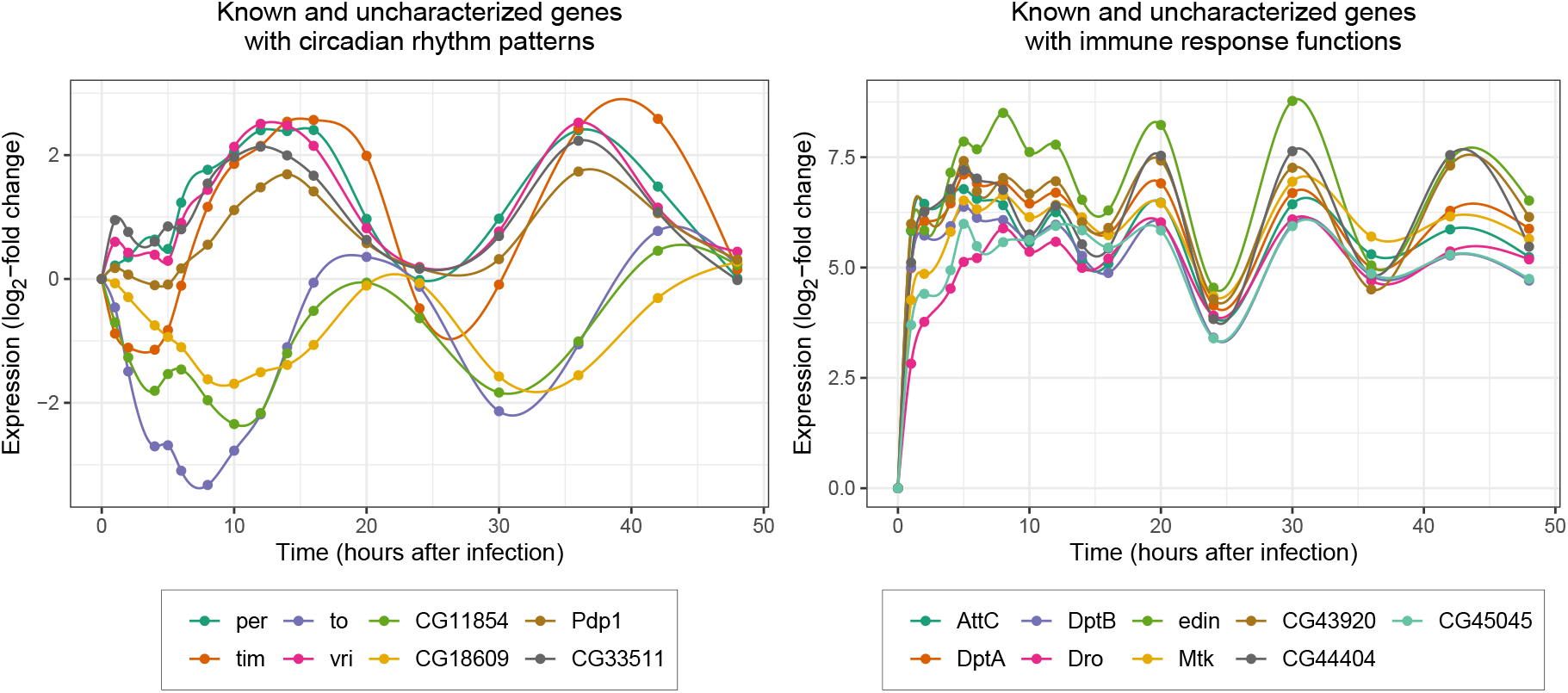
Left: The uncharacterized gene CG33511 exhibits similar time dynamics to known circadian rhythm genes with 24-hour cyclic temporal expressions. Right: Uncharacterized genes CG43920 and CG45045 exhibit similar temporal expression patterns to known immune response genes, which are up-regulated in response to infection.

Incorporating pathway membership or protein-protein interaction networks into the lead-lag *R*^2^ enables us to encourage genes in the same pathways to receive higher pairwise similarity scores, thus separating true positives from false positives. Importantly, we can also suggest possible pathways for previously uncharacterized genes. In our method, external biological information is used to determine the locations of normal prior distributions on the parameters *c*_1_, …, *c*_5_ in the model (4). Upon obtaining the posterior estimates of these parameters, we recompute the lead-lag *R*^2^ to obtain the *Bayesian lead-lag R*^2^ (LL*R*^2^). In doing so, we observe a desirable shrinkage effect in the distribution of the Bayesian lead-lag *R*^2^ values that pares down the number of significant associations. We next detail the hierarchical model and its hyperparameters in Section 3.

## 3 Empirical Bayes methodology

In this section, we propose our empirical Bayes approach to deriving biologically-informed similarity metrics between genes for clustering and network analysis. The components of our method are: 1) encoding external biological information into a prior adjacency matrix, 2) defining a normal-inverse gamma prior, specifically Zellner’s *g*-prior, on the parameters of the ODE-based model of gene expression (4), 3) optimally selecting the hyperparameter *g* in Zellner’s *g*-prior, and 4) calculating the Bayesian lead-lag *R*^2^. Note that parts 2-4 lead to the computation of the Bayesian lead-lag *R*^2^ for a single gene pair; a summary of the full algorithm for all pairs is provided in Appendix B.

### 3.1 Part 1: Leveraging biological priors

There exist numerous databases that extensively document known or predicted interactions between genes as well as their functional roles. For instance, Gene Ontology (GO) terms are keywords that describe a gene’s molecular function, role in a biological process (e.g., “lipid metabolism”), or cellular localization (e.g., “nucleus”) [Ashburner et al., 2000]. Semantic similarity methods, such as the R package `GOSemSim` [Yu et al., 2010], have been developed to determine how related genes are based on their associated GO terms. Other curated databases that similarly assign genes to pathways include KEGG [Kanehisa and Goto, 2000], Reactome [Fabregat et al., 2018], and BioCyc [Karp et al., 2019]. The STRING database [Szklarczyk et al., 2019] aggregates multiple sources of information to generate a more holistic measurement of the association between two genes. For each pair of genes in an organism, STRING provides a score between 0 and 1 indicating how likely the two genes’ encoded proteins are to interact physically based on experimental evidence, belong to the same pathways according to the aforementioned databases, be conserved across species, or be mentioned in the same studies.

Regardless of which sources of external biological information one employs, the first step of our method is to encode this information into a matrix **W** of size *N* × *N*, where *N* is the total number of genes under study. The entries of **W** are:

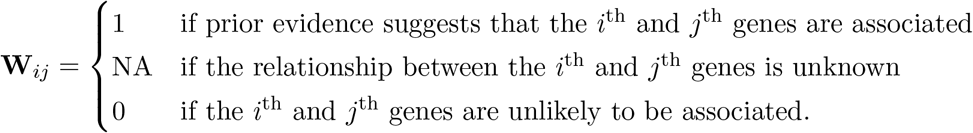

Intuitively, **W** can be viewed as an adjacency matrix for a network that reflects the known relationships amongst the genes. Our proposed method uses this network as an informed basis for the network constructed from the time-course gene expression data itself. Importantly, the data can indicate what the “NA” entries of **W** ought to be, as well as confirm (or refute) the “0” or “1” entries.

In the event that **W** consists largely of “NA” entries, one can also use time-course gene expression data from previous studies to fill in some of them. Such datasets often consist of multiple replicates of the expression measurements, possibly gathered under the same experimental conditions to account for sampling variation, or different conditions to assess the effect of a treatment.

### 3.2 Part 2: The normal-inverse gamma model and Zellner’s *g*-prior

Recall from Section 2.1 that given measurements {*m*_*A*_(*t*_1_), …, *m*_*A*_(*t*_*n*_)} and {*m*_*B*_(*t*_1_), …, *m*_*B*_(*t*_*n*_)} of the expressions of two genes *A* and *B* at times *t*_1_, …, *t*_*n*_, the model we aim to fit is

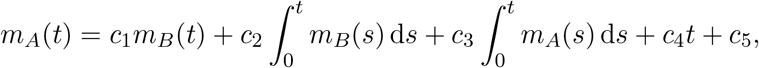

where *c*_1_, …, *c*_5_ are unknown parameters, and the integrals 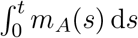 and 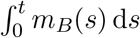 are estimated by numerically integrating spline interpolants of the given data. As seen in (5), this model can be represented in matrix form as **Y** = **X*β*** + ***ε***, where ***ε*** ∼ *N* (**0**, *σ*^2^**I**_*n*_).

Since *c*_1_ and *c*_2_ link the expressions of genes *A* and *B*, we intuitively ought to place priors of non-zero mean on these two parameters if **W** indicates that the genes are associated. To do this, we adopt the normal-inverse gamma model for ***β*** and *σ*^2^, which is used frequently in Bayesian regression and allows for flexible modeling of the prior mean and covariance matrix of ***β***. The normal-inverse gamma model specifies ***β***|*σ*^2^ ∼ *N* (***β***_0_, *σ*^2^**V**_0_) and *σ*^2^ ∼ Γ^−1^(*a, b*) where ***β***_0_ ∈ ℝ^*p*×1^, **V**_0_ ∈ ℝ^*p*×*p*^ is a positive semidefinite matrix, and *a, b* > 0. It is then said that (***β***, *σ*^2^) jointly follow a normal-inverse gamma distribution with parameters (***β***_0_, **V**_0_, *a, b*). This is a conjugate prior, so the posterior distribution of (***β***, *σ*^2^) is also normal-inverse gamma with parameters (***β***_*_, **V**_*_, *a*_*_, *b*_*_) defined as

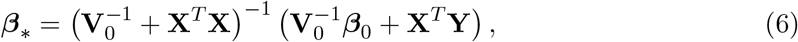

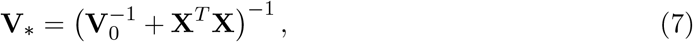

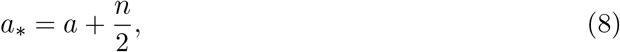

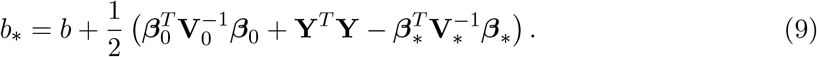

That is, the conditional posterior of ***β*** given *σ*^2^ and the posterior of *σ*^2^ are ***β***|*σ*^2^, **Y** ∼ *N* (***β***_*_, *σ*^2^**V**_*_) and *σ*^2^|**Y** ∼ Γ^−1^(*a*_*_, *b*_*_). The posterior mean ***β***_*_ can be used as the estimated regression parameters. This estimator has the desirable properties of consistency and asymptotic efficiency in large samples and admissibility in finite samples [Giles and Rayner, 1979].

The hyperparameters ***β***_0_ and **V**_0_ are of particular interest as they allow us to incorporate biological information into our model. In defining ***β***_0_, recall that we wish to place priors with non-zero mean on the parameters *c*_1_ and *c*_2_ when external sources suggest that genes *A* and *B* are co-regulated or at least associated. We noted in Section 2.1 that *c*_1_ represents the ratio *α*_*A*_*/α*_*B*_, where *α*_*A*_ and *α*_*B*_ denote the strength of a common regulatory signal in the first-order dynamics of the two genes. If the genes are associated, it is reasonable to believe *a priori* that *α*_*A*_ = *α*_*B*_, implying *c*_1_ = 1. Then we could also say *a priori* that *c*_2_ = 1, since *c*_2_ represents a parameter that is proportional to *c*_1_. When prior information about genes *A* and *B* is lacking or suggests that they are unrelated, a prior mean of zero for *c*_1_ and *c*_2_ is appropriate. Supposing genes *A* and *B* are the *i*^th^ and *j*^th^ genes in the dataset, we can thus set the prior mean ***β***_0_ as follows:

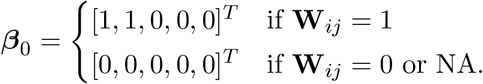

As for the prior covariance matrix *σ*^2^**V**_0_ of ***β***, we first note that for the linear model **Y** = **X*β*** + ***ε*** where ***ε*** has the covariance matrix *σ*^2^**I**_*n*_, the least-squares estimator 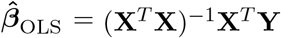 has the covariance matrix *σ*^2^(**X**^*T*^ **X**)^−1^. A popular choice for **V**_0_ is therefore *g*(**X**^*T*^ **X**)^−1^, where *g* > 0. This choice of **V**_0_ yields a particularly tractable special case of the normal-inverse gamma model known as *Zellner’s g-prior* [Zellner, 1986]. Substituting this choice of **V**_0_ into the posterior mean ***β***_*_ in (6) and covariance matrix **V**_*_ in (7), we obtain

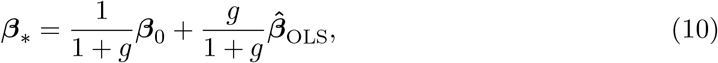

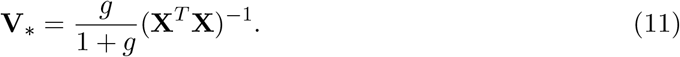

(10) reveals that under Zellner’s *g*-prior, the posterior mean ***β***_*_ is a convex combination of the prior mean ***β***_0_ and the least-squares estimator 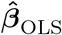. The parameter *g* balances the weights placed on external information encoded by ***β***_0_ and on the data used to compute 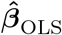, so the selection of *g* is an important component of our modeling strategy. We next describe our data-driven approach to choosing it.

### 3.3 Part 3: Optimal selection of *g* in Zellner’s *g*-prior

Several methods for choosing *g* in Zellner’s *g*-prior have been proposed previously. For instance, George and Foster [2000] discuss an empirical Bayes method in which one selects the value of *g* that maximizes the marginal likelihood of **Y**. Liang et al. [2008] provide a closed form expression for this maximizing value of *g* that is nearly identical to the *F*-statistic for testing the hypothesis that ***β*** = **0**. They show that this maximum marginal likelihood approach has the desirable property that the Bayes factor for comparing the full model to the null (intercept-only) model diverges as the *R*^2^ approaches 1. For our application, one concern is that the *F*-statistic defining this particular estimate of *g* is likely to be overwhelmingly large for many gene pairs. If *g* is large, then ***β***_*_ will be very close to 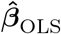 according to (10). As a result, any biological evidence of association captured by ***β***_0_ will have a negligible impact on the model.

A fully Bayesian approach to selecting *g* involves placing a prior distribution on *g* that is then integrated out when defining the prior on ***β***. This method is motivated by Zellner and Siow [1980], who propose placing a Cauchy prior on ***β*** to derive a closed-form posterior odds ratio for hypothesis testing purposes. These priors can be represented as a mixture of *g*-priors with an inverse gamma prior on *g*, although a closed-form posterior estimate of *g* is unavailable. Cui and George [2008] and Liang et al. [2008] alternatively consider a class of priors of the form 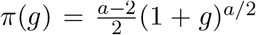 for *a* > 2, known as the hyper-*g* priors. Under these priors, the posterior mean of *g* can be expressed in closed form in terms of the Gaussian hypergeometric function.

Because our application involves fitting regression models for potentially thousands of gene pairs, the computational cost of fully Bayesian methods for selecting *g* requires us to consider alternative approaches. One idea is to select the value of *g* that minimizes the sum of squared residuals ‖**Y**− **Ŷ**‖^2^, where **Ŷ** = **X*β***_**\***_ is the vector of fitted values:

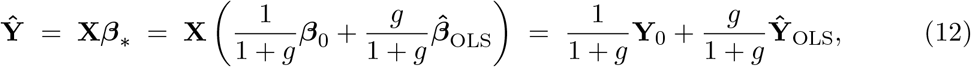

where **Y**_0_ = **X*β***_0_ and 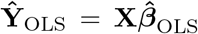. However, we found that there are no analytical solutions to *g*_*_ = argmin_*g*>0_ ‖**Y −Ŷ**‖^2^ = 0. Instead, we can minimize Stein’s unbiased risk estimate (SURE), which is an unbiased estimate of ‖**Ŷ − X*β***‖^2^. Below is a rephrased version of Theorem 8.7 in Fourdrinier et al. [2018], which defines SURE for the general problem of estimating 𝔼 (**Y**) using a linear estimator of the form **Ŷ** = **a** + **SY**. This theorem statement differs from its original version in that we have rewritten the divergence of 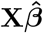 with respect to **Y** using the generalized degrees of freedom developed by Efron [2004].

#### Theorem 3.1

(SURE for linear models). Let **Y** ∼ *N* (**X*β***, *σ*^2^**I**_*n*_), where the dimensions of **X** are *n* × *p*, and let 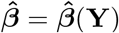 be a weakly differentiable function of the least squares estimator 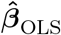 such that 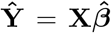 can be written in the form **Ŷ = a** + **SY** for some vector **a** and matrix **S**. Let 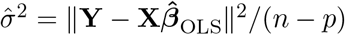. Then,

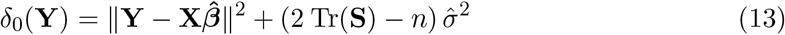

is an unbiased estimator of ‖**Ŷ−X*β***‖^2^.

From (12), observe that we can write **Ŷ** as 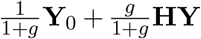, where **H** = **X**(**X**^*T*^ **X**)^−1^**X**^*T*^. Therefore the matrix **S** in Theorem 3.1 is *g***H***/*(1 + *g*), whose trace is *gp/*(1 + *g*). SURE in (13) then becomes

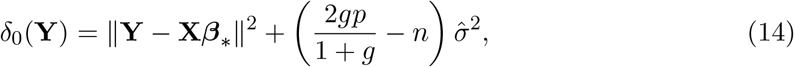

where we have also substituted the posterior mean ***β***_*_ in (10) for 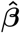.

We next present the value of *g* that minimizes SURE.

#### Theorem 3.2

(SURE minimization with respect to *g*). The value of *g* that minimizes SURE in (14) is

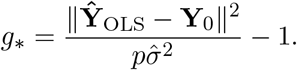

The proof of Theorem 3.2 is provided in Appendix A.

It is quite possible that 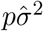 is small, resulting in a large value of *g*_*_ (i.e., *g*_*_ ≫ 1) in Theorem 3.2. In this case, ***β***_*_ in (10) will be largely influenced by the data via 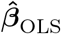, rather than by prior information via ***β***_0_. This is desirable when the relationship between the two genes is unknown (i.e. **W**_*ij*_ = NA), but not if the relationship is known to be unlikely (i.e. **W**_*ij*_ = 0). In the latter case, we prefer to shrink the regression coefficients towards the prior mean ***β***_0_ = **0** so as to yield a smaller lead-lag *R*^2^ value. To address this, we set *g* conditionally on **W**_*ij*_ as

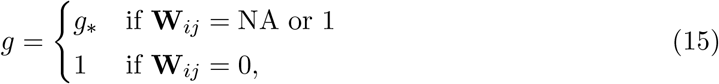

where *g*_*_ is defined according to Theorem 3.2.

### 3.4 Part 4: Computing the *R*^2^ for Bayesian regression models

Once a posterior estimate of the model coefficients (10) has been obtained, with the parameter *g* selected optimally, we can compute the Bayesian lead-lag *R*^2^ between genes *A* and *B*.

Recall that for a linear model **Y** = **X*β*** + ***ε*** where ***β*** is estimated by the least-squares estimator 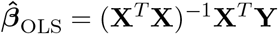, the *R*^2^ is defined as

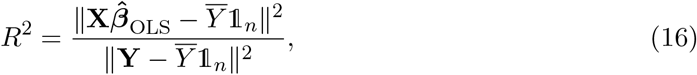

where 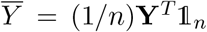. In Bayesian regression, however, the standard decomposition of total sum of squares into residual and regression sums of squares no longer holds. Thus, when we replace 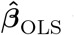 with an estimator such as the posterior mean of ***β***, the formula (16) can potentially yield *R*^2^ > 1. As a remedy to this issue, we compute the *R*^2^ as the ratio of the variance of the model fit to itself plus the variance of the residuals. This ratio is within [0, 1] by construction, and is given by

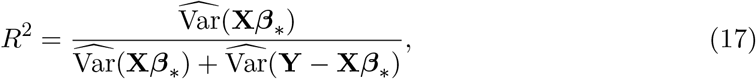

where, for a vector **Z** = [*z*_1_ … *z*_*n*_]^*T*^ with mean 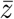, we define 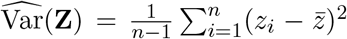. This calculation is based on the approach to computing *R*^2^ for Bayesian regression models proposed in Gelman et al. [2018]. For our application, we refer to (17) as the *Bayesian lead-lag R*^2^ (LL*R*^2^).

### 3.5 Clustering and empirical analysis with the Bayesian lead-lag *R*^2^

Given a dataset of *N* genes whose expressions are measured at *n* time points, our objective is to cluster the genes based on their temporal expression patterns. To do this, we compute a *N* × *N* similarity matrix **S** where **S**_*i,j*_ stores the Bayesian LL*R*^2^ in (17) between the *i*^th^ and *j*^th^ genes. We then apply hierarchical clustering to the distance matrix **J** − **S**, where **J** is the *N* × *N* matrix of ones.

Note that the Bayesian LL*R*^2^ is asymmetric: LL*R*^2^(*i, j*) ≠ LL*R*^2^(*j, i*). Here, LL*R*^2^(*i, j*) denotes the Bayesian LL*R*^2^ where we treat the *i*^th^ gene as the response (gene *A*) in the model (4). For our purpose of clustering a large set of genes for empirical analysis, we symmetrize the Bayesian LL*R*^2^ by setting:

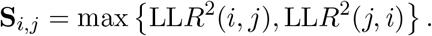

In practice, it is also common to use similarity measures such as the Bayesian LL*R*^2^ to produce a ranked list of gene-gene associations. To aid this procedure, we further propose two additional metrics that one could use in conjunction with the Bayesian LL*R*^2^. These metrics are derived from the following two sub-models of the original model (4):

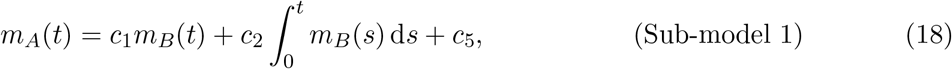

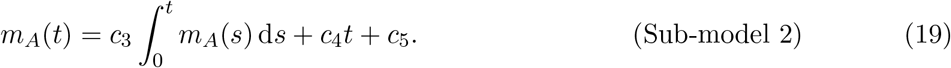

The first sub-model describes the temporal expression of gene *A* as a function only of the potentially co-regulated gene *B*. The second sub-model is “autoregressive” in the sense that it describes gene *A*’s expression only in terms of its own past and linear time trends. We again apply our Bayesian approach to fitting these two sub-models and compute new variants of the Bayesian LL*R*^2^ from each, denoted 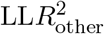 and 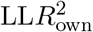 respectively. The 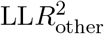 value indicates the amount of variation in gene *A*’s temporal expression that can be explained by the dynamics of an*other* gene *B*. 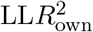 indicates the amount of variation in gene *A* that is explained by its *own* past, via its time integrals, and linear time trends. We can view 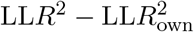 as the amount of *additional* variation in gene *A*’s temporal dynamics that can be explained by considering gene *B*, on top of the variation captured by gene *A*’s own past via sub-model 2. Intuitively, if LL*R*^2^ is large, then a large value of 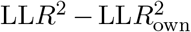 suggests that the LL*R*^2^ value is unlikely to mean a false positive relationship between the two genes. In Section 4.4, we demonstrate empirically that using 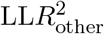 and 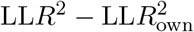 together can help identify gene pairs with highly similar time dynamics.

### 3.6 Significance of the Bayesian lead-lag *R*^2^

One may further desire a notion of statistical significance for the Bayesian LL*R*^2^. One option is to simulate the posterior distribution of the LL*R*^2^ using draws from the posterior distribution of ***β***, as described in Gelman et al. [2018]. Recall from Section 3.2 that the posterior distribution of (***β***, *σ*^2^) is normal-inverse gamma with parameters (***β***_*_, **V**_*_, *a*_*_, *b*_*_), defined respectively in (10), (11), (8), and (9). To draw samples from this posterior distribution, we can first sample *σ*^2^ from the Γ^−1^(*a*_*_, *b*_*_) distribution, and then sample ***β*** from the *N* (***β***_*_, *gσ*^2^**V**_*_) distribution. In particular, if **W**_*ij*_ = 1, we may wish to simulate the posterior distribution of the Bayesian LL*R*^2^ under a null hypothesis of no relationship between genes *A* and *B*. This can be reflected in the sampling procedure by calculating ***β***_*_ with ***β***_0_ set to **0**.

Alternatively, one could use the Bayes factors presented in Liang et al. [2008] to corroborate the Bayesian lead-lag *R*^2^. Let ℳ _*F*_ denote the model (4) of gene expression, which has an intercept and *p* = 4 “covariates” with coefficients *c*_1_ through *c*_4_. Let ℳ _*N*_ denote the null (intercept-only) model. Then the Bayes factor for comparing ℳ _*F*_ to ℳ _*N*_ is

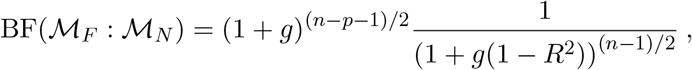

where *R*^2^ is the usual coefficient of determination in (16). Kass and Raftery [1995] interpret a log_10_(BF(ℳ _*F*_ : ℳ _*N*_)) value between 1 and 2 as “strong” evidence in favor of ℳ _*F*_, or above 2 as “decisive” evidence.

In Appendix B, we give a summary of our methodology in the form of a generic algorithm that can be run on any time-course gene expression dataset.

### 3.7 Possible modifications to prior hyperparameters

In Section 3.2, we set the prior mean ***β***_0_ of the parameters of the ODE model to [1, 1, 0, 0, 0]^*T*^ when **W**_*ij*_ = 1, i.e. there is prior evidence suggesting genes *A* and *B* are associated. To make the method even more data-driven, one could alternatively set ***β***_0_ = [*ξ, ξ*, 0, 0, 0]^*T*^ in the **W**_*ij*_ = 1 case, where *ξ* ≠ 0 is chosen adaptively from the data. The following theorem presents the values of *ξ* and *g* that simultaneously minimize SURE in (14) in this setting.

#### Theorem 3.3

(SURE minimization with respect to *ξ* and *g*). Assume the entries of the least-squares estimator 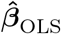 are all distinct and the expression of gene *B* is non-zero for at least one time point, i.e. *m*_*B*_(*t*_*i*_) ≠ 0 for at least one *i*. Let ***β***_0_ = [*ξ, ξ*, 0, 0, 0]^*T*^ in the case that **W**_*ij*_ = 1. Then the values of *ξ* and *g* that minimize SURE in (14) are

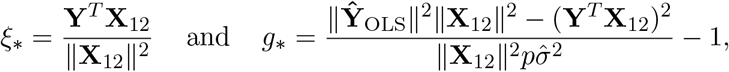

where **X**_12_ ∈ ℝ ^*n*×1^ is the element-wise sum of the first two columns of **X**. The proof of Theorem 3.3 is provided in Appendix A.

## 4 Results for the *Drosophila melanogaster* dataset

### 4.1 Collecting a time-course gene expression dataset

The expression of a gene is typically measured via RNA sequencing (RNA-seq) as the count of a particular type of messenger RNA found in a tissue sample. These counts can be normalized to library size and transformed using the limma-voom transformation [Law et al., 2014]. This transformation produces near-normally distributed gene expression measurements, making them more amenable to analysis with linear models such as those described in Section 2.1.

Our primary time-course gene expression dataset, introduced in Schlamp et al. [2021], profiles the expression dynamics of 12,657 genes in *Drosophila melanogaster* (fruit fly) in response to an immune challenge. Immune responses were triggered in flies by injecting them with commercial lipopolysaccharide, which contains peptidoglycan (cell wall molecules) derived from the bacterium *E. coli*. Following injection, the flies were sampled via RNA-seq at 20 time points over five days. The data was normalized by the aforementioned limma-voom transformation and expressed as the log_2_-fold change with respect to the first time point, which was sampled pre-injection as a control. We focus on the first 17 time points, ranging from zero to 48 hours post-injection, as this is when most of the variation in expression occurs.

Differential expression analysis is typically used to identify genes exhibiting significant expression changes, and thus reduce a set of thousands of genes into a smaller set meriting further study. In Appendix C, we provide further details on how we use such methods to reduce our set of 12,657 genes into a set of 1735 genes of interest. We also describe therein how we define our prior adjacency matrix **W**, introduced in Section 3.1, using the STRING database.

### 4.2 Small-scale case study: immunity and metabolism

We first validate our methodology on a small set of genes whose behavior exhibits a known interplay between immunity and metabolism. Schlamp et al. [2021] observe that exposure to bacterial peptidoglycan has an effect not only on the time dynamics of immune response, but also on the expression of genes involved in metabolism. In particular, some genes involved in immune response are up-regulated immediately following peptidoglycan injection, while other genes associated with metabolic processes are down-regulated more slowly. Interestingly, the metabolic genes return to their pre-infection levels of expression well before the immune response has subsided. This phenomenon can be observed in Figure 4, which shows the expression patterns of two immune response genes (*IM1, IM2*) and four metabolic process genes (*FASN1, UGP, mino, fbp*).

**Figure 4:**
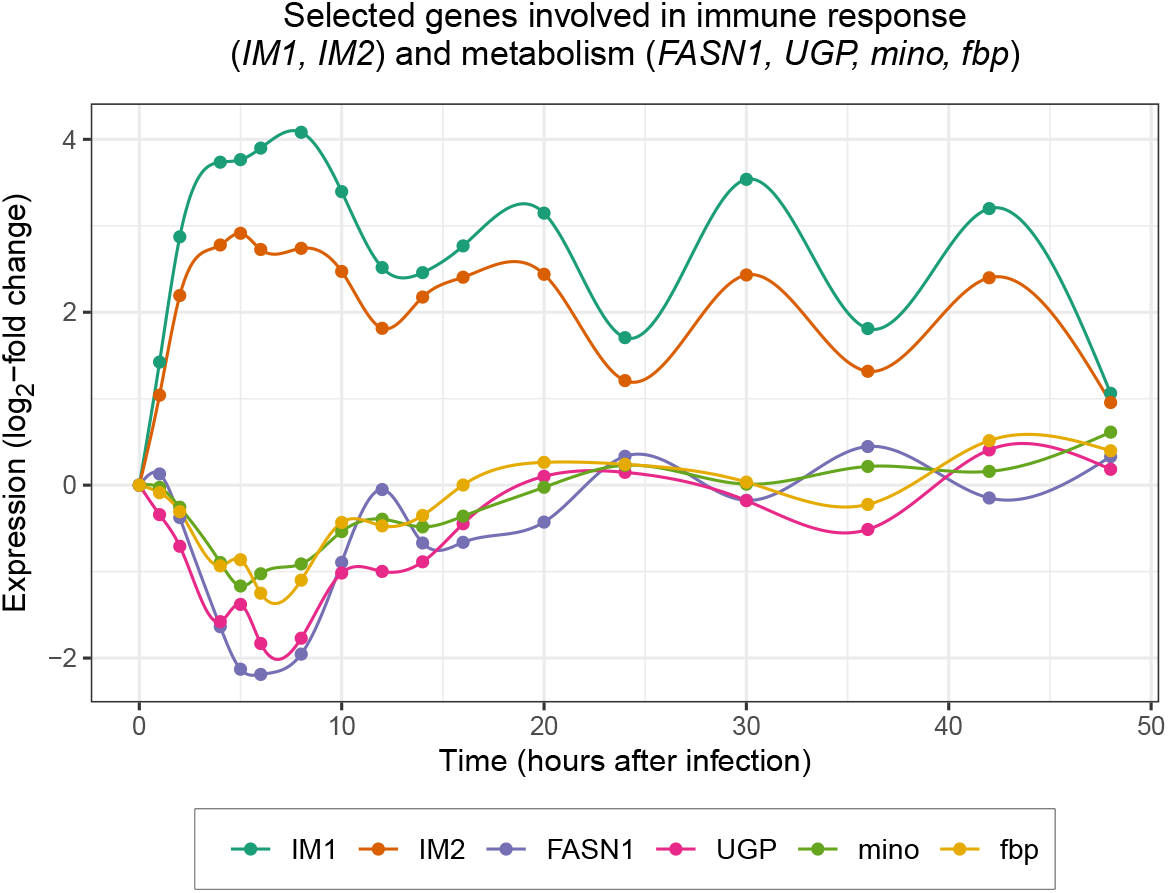
Temporal expression patterns of selected genes involved in immune response (IM1, IM2) and metabolic processes (FASN1, UGP, mino, fbp). Upon infection, immune response genes are immediately up-regulated. At the same time, metabolic genes are down-regulated but return to pre-infection expression levels after 12 to 24 hours, which is before the immune response is resolved.

Tables 1-3 show the prior adjacency matrix **W**, the Bayesian LL*R*^2^ values, and the non-Bayesian LL*R*^2^ values (computed via ordinary least-squares regression) corresponding to these six genes.

**Table 1:** *Portion of the prior adjacency matrix* **W** *corresponding to the six genes in Figure 4. Red cells indicate that there is prior evidence of association between the two genes. “NA” entries indicate that the association between the two genes is unavailable in the STRING database*.

|  | IM1 | IM2 | FASN1 | UGP | mino | fbp |
| --- | --- | --- | --- | --- | --- | --- |
| IM1 | - | 1 | NA | 0 | 0 | NA |
| IM2 | 1 | - | NA | 0 | 0 | NA |
| FASN1 | NA | NA | - | NA | NA | NA |
| UGP | 0 | 0 | NA | - | 1 | NA |
| mino | 0 | 0 | NA | 1 | - | NA |
| fbp | NA | NA | NA | NA | NA | - |

Within this set of six genes, Table 1 indicates that there is prior evidence of association between the immune response genes *IM1* and *IM2*, as well as between the metabolic genes *mino* and *UGP*. The off-diagonal “NA” entries in Table 1 signify that the relationships between the immune and metabolic genes are uncharacterized in the STRING database. However, the Bayesian LL*R*^2^ values between the metabolic gene *FASN1* and both immune response genes are high, as shown in Table 2, indicating that the relationship identified by Schlamp et al. [2021] between these gene groups is automatically uncovered by our proposed Bayesian method. Indeed, Figure 4 shows that the temporal expression pattern of *FASN1* resembles a vertically-reflected copy of that of either *IM1* or *IM2*. Table 3 shows the non-Bayesian LL*R*^2^ values for each gene pair and demonstrates that computing the LL*R*^2^ without biologically-informed priors yields inflated scores that make it difficult to discern either within- or between-group associations. For further validation of these results, we present Tables 4 and 5 in Appendix E. These tables display the values of the metric 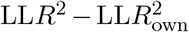,which we introduced in Section 3.5 as a means of assessing whether associations indicated by the LL*R*^2^ alone are false positives.

**Table 2:** Bayesian LLR^2^ values corresponding to each gene pair in Figure 4. Colored cells mark values above 0.76, the 95^th^ percentile of the empirical distribution of this metric for this dataset. Red cells are consistent with prior evidence of association (c.f. red cells in Table 1). Blue cells indicate potential associations that may not have been known previously. Overall, colored cells point to biologically-meaningful relationships within the immunity and metabolism gene groups as well as between these groups. Between-group association is suggested by the high LLR^2^ values between FASN1 and both IM1 and IM2. Indeed, Figure 4 shows that the expression pattern of FASN1 is a vertical reflection of these immune response genes’ patterns.

|  | IM1 | IM2 | FASN1 | UGP | mino | fbp |
| --- | --- | --- | --- | --- | --- | --- |
| IM1 | - | 0.99 | 0.82 | 0.30 | 0.40 | 0.68 |
| IM2 | 0.99 | - | 0.80 | 0.30 | 0.39 | 0.66 |
| FASN1 | 0.82 | 0.80 | - | 0.83 | 0.98 | 0.82 |
| UGP | 0.30 | 0.30 | 0.83 | - | 0.91 | 0.99 |
| mino | 0.40 | 0.39 | 0.98 | 0.91 | - | 0.90 |
| fbp | 0.68 | 0.66 | 0.82 | 0.99 | 0.90 | - |

**Table 3:** Non-Bayesian LLR^2^ values corresponding to each gene pair in Figure 4. Colored cells mark values above 0.96, the 95^th^ percentile of the empirical distribution of this metric for this dataset. Colors have the same interpretation as in Table 2. Without the proposed Bayesian method, within-group and between-group associations are not identified as clearly as in Table 2. Furthermore, nearly all values in the table are close to the selected threshold of 0.96, suggesting that LLR^2^ values are easily inflated without the use of priors, even for dissimilar temporal expression patterns (such as those of mino and IM1).

|  | IM1 | IM2 | FASN1 | UGP | mino | fbp |
| --- | --- | --- | --- | --- | --- | --- |
| IM1 | - | 1.00 | 0.97 | 0.94 | 0.97 | 0.93 |
| IM2 | 1.00 | - | 0.96 | 0.94 | 0.97 | 0.93 |
| FASN1 | 0.97 | 0.96 | - | 0.91 | 0.98 | 0.88 |
| UGP | 0.94 | 0.94 | 0.91 | - | 0.93 | 0.99 |
| mino | 0.97 | 0.97 | 0.98 | 0.93 | - | 0.93 |
| fbp | 0.93 | 0.93 | 0.88 | 0.99 | 0.93 | - |

**Table 4:** Values of the Bayesian 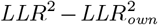 corresponding to each gene pair in Figure 4. Colored cells mark values above 0.20, the 95^th^ percentile of the empirical distribution of this metric for this dataset. Red cells are consistent with prior evidence of association, and blue cells point to potential associations that may not have been known previously. Most of the colored cells in Table 2 are colored here as well, indicating that the associations drawn from Table 2 are unlikely to be false positives.

|  | IM1 | IM2 | FASN1 | UGP | mino | fbp |
| --- | --- | --- | --- | --- | --- | --- |
| IM1 | - | 0.64 | 0.27 | 0.05 | 0.09 | 0.09 |
| IM2 | 0.64 | - | 0.26 | 0.04 | 0.09 | 0.06 |
| FASN1 | 0.27 | 0.26 | - | 0.20 | 0.27 | 0.23 |
| UGP | 0.05 | 0.04 | 0.20 | - | 0.20 | 0.39 |
| mino | 0.09 | 0.09 | 0.27 | 0.20 | - | 0.19 |
| fbp | 0.09 | 0.06 | 0.23 | 0.39 | 0.19 | - |

**Table 5:** Values of the non-Bayesian 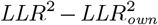 corresponding to each gene pair in Figure 4. Colored cells mark values 0.13, the 95^th^ percentile of the empirical distribution of this metric for this dataset. Colors have the same interpretation as in Table 4. Most of the colored cells in Table 3, which displays the non-Bayesian LLR^2^ values, are not highlighted in this table; thus, the non-Bayesian 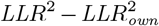 metric is less helpful than its Bayesian counterpart in verifying whether or not the associations suggested by the LLR^2^ alone are false positives.

|  | IM1 | IM2 | FASN1 | UGP | mino | fbp |
| --- | --- | --- | --- | --- | --- | --- |
| IM1 | - | 0.09 | 0.06 | 0.03 | 0.06 | 0.02 |
| IM2 | 0.09 | - | 0.06 | 0.03 | 0.06 | 0.02 |
| FASN1 | 0.06 | 0.06 | - | 0.09 | 0.17 | 0.13 |
| UGP | 0.03 | 0.03 | 0.09 | - | 0.11 | 0.17 |
| mino | 0.06 | 0.06 | 0.17 | 0.11 | - | 0.11 |
| fbp | 0.02 | 0.02 | 0.13 | 0.17 | 0.11 | - |

### 4.3 Biologically-meaningful clustering with the Bayesian LL*R*^2^

We now apply hierarchical clustering using Ward’s method [Ward Jr, 1963] to the *N* × *N* distance matrix **J** − **S**, where **J** is a matrix of ones and **S**_*i,j*_ = max{LL*R*^2^(*i, j*), LL*R*^2^(*j, i*)} is the similarity matrix containing the symmetrized Bayesian LL*R*^2^ values between all gene pairs. These genes come from the dataset described in Section 4.1, so *N* = 1735. Cutting the resulting dendrogram at a height of ten yields 12 clusters, which we display in Figure 5.

**Figure 5:**
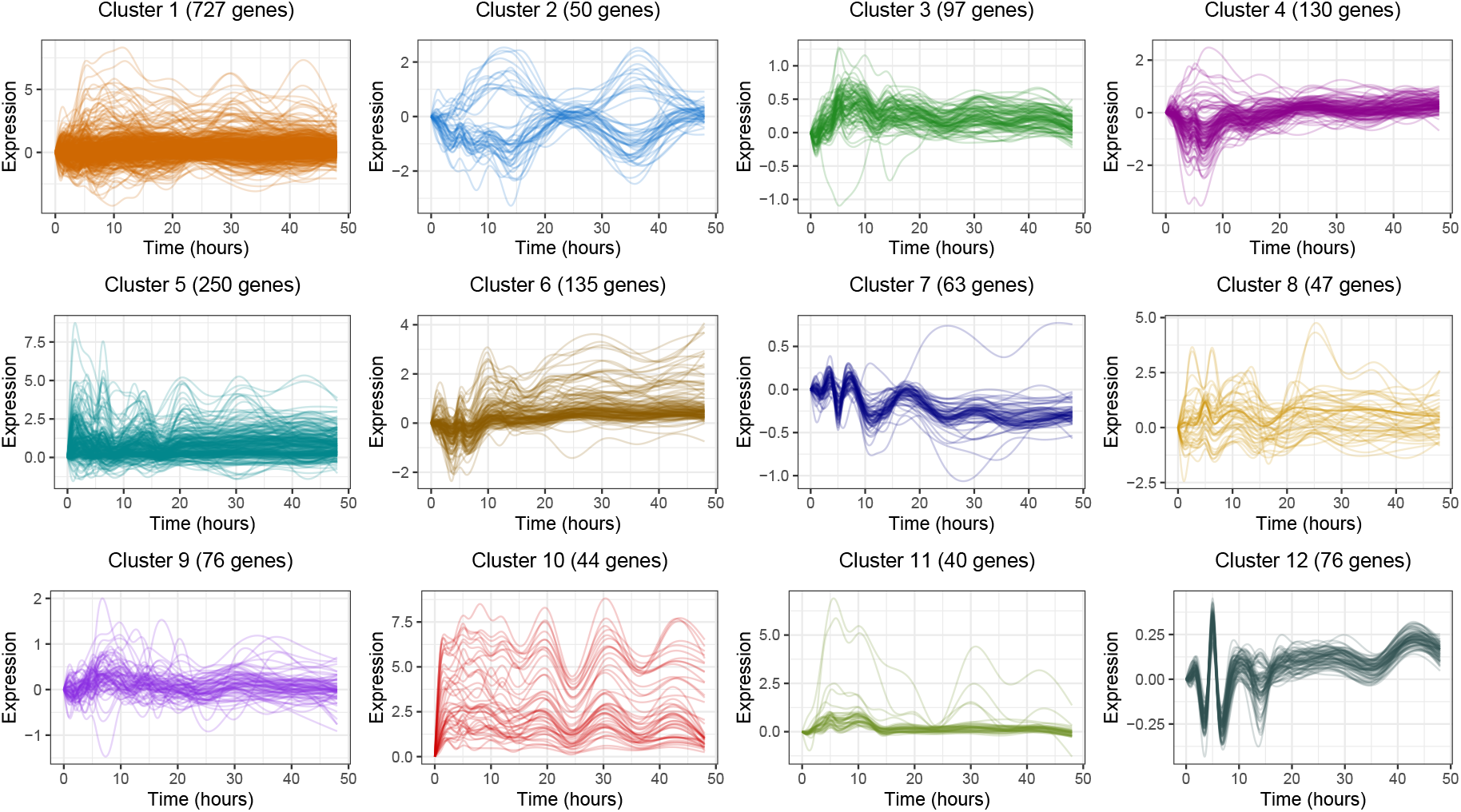
*Plots of the temporal expression patterns of genes in each cluster obtained by applying hierarchical clustering to the distance matrix* **J** − **S**, *where* **S** *contains the Bayesian LLR*^2^ *for all gene pairs. Cluster sizes range from 40 to 727 genes, with a mean of 145 and a median of 76. These plots visually demonstrate that the Bayesian LLR*^2^ *is capable of capturing similarities in the overall shapes of two temporal expression patterns, even if one gene is a time-delayed copy of the other or is reflected across a horizontal axis, for instance*.

From each cluster, we can also construct a network in which an edge is defined between two genes if their corresponding Bayesian LL*R*^2^ exceeds a certain threshold. In our analysis, we choose this threshold to be 0.9, which is also the 99^th^ percentile of the empirical distribution of the Bayesian LL*R*^2^ for this dataset.

To understand the biological processes that are represented by these clusters, we make use of the Gene Ontology (GO) resource [Ashburner et al., 2000, Carbon et al., 2021]. GO links genes with standardized terms that describe their functions. To determine whether a GO term is significantly enriched within a cluster, i.e. whether the cluster contains significantly more genes associated with that term than expected by chance, we perform a hypergeometric test using the R package `ClusterProfiler` [Yu et al., 2012]. We use Benjamini-Hochberg (B-H) corrected *p*-values below 0.05 [Benjamini and Hochberg, 1995] from this test to determine “significant” enrichment.

Our analysis shows that with the exception of cluster 8, all clusters are significantly enriched for specific biological functions. Recall that our dataset profiles the dynamics of immune response, which is an energetically costly process that is also associated with metabolic changes [DiAngelo et al., 2009]. The interplay between immunity and metabolism, which we briefly explored in Section 4.2, is represented particularly well in these clusters. Clusters 1, 4, 6, and 7 are significantly enriched for metabolic processes; cluster 10 is significantly enriched for immune response; and cluster 5 is significantly enriched for both metabolic processes and immune responses. Below, we highlight biologically relevant findings from one cluster, and we discuss three additional clusters in Appendix F. These examples show that clustering with the Bayesian LL*R*^2^ allows genes with similar but lagged expression patterns to be grouped together, even in the absence of known prior information. Finally, the Bayesian LL*R*^2^ is not influenced by the direction of gene expression changes (i.e., positive or negative changes), making it easier to detect tradeoffs or negative regulatory interactions between genes. As a comparison, Figures 11 and 12 in Appendix D present clustering results obtained using the non-Bayesian LL*R*^2^ as well as Pearson correlation between genes, though neither method groups together genes with similarly-shaped trajectories as effectively.

#### 4.3.1 Analysis of cluster 10

In cluster 10, which consists of 44 genes, one of the most significantly enriched GO terms is “defense response”. This GO term is supported by 24 genes in the cluster, within which there are two distinct groups that we display in Figure 6: eight genes that are known to respond to “Imd” signaling and eight genes that respond to “Toll” signaling. The Imd and Toll signaling pathways represent well-studied molecular responses to infection in flies. The Imd pathway is tailored to fight off infections from gram-negative bacteria [Kaneko et al., 2004, Zaidman-Rémy et al., 2011, Hanson and Lemaitre, 2020], while the Toll pathway fights off infections from gram-positive bacteria and fungi [Gobert et al., 2003, Hanson and Lemaitre, 2020].

**Figure 6:**
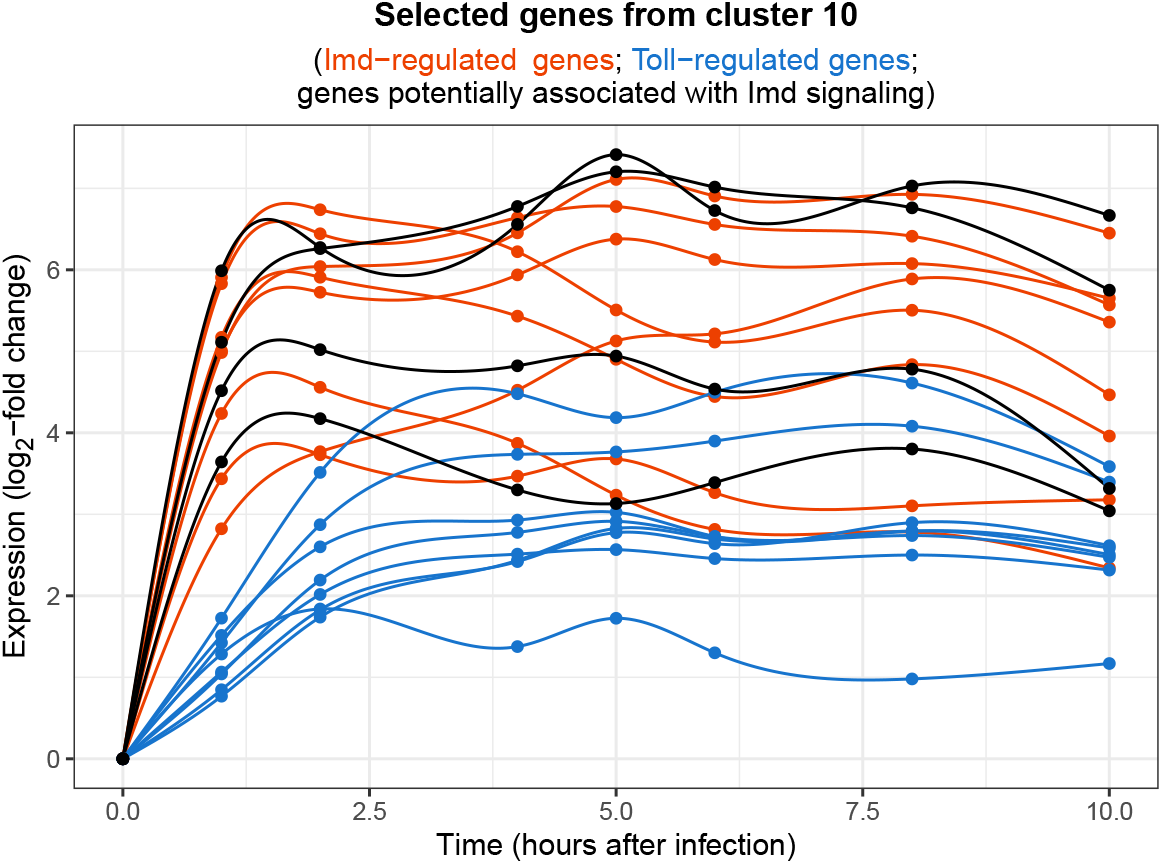
Temporal expression patterns of selected genes in cluster 10 during the first ten hours following peptidoglycan injection. The eight red lines correspond to Imd-regulated genes. The eight blue lines correspond to Toll-regulated genes, which show a smaller and delayed up-regulation. The four black lines correspond to genes that exhibited high Bayesian LLR^2^ values (> 0.9) with Imd-regulated genes; while there is no prior information in STRING linking them with the Imd pathway, their co-clustering with Imd-regulated genes was also observed in Schlamp et al. [2021].

Since the flies profiled in this gene expression dataset were injected with peptidoglycan derived from *E. coli*, a gram-negative bacterium, we expect to see an activation of Imd-regulated genes. Indeed, as seen in Figure 6, the eight Imd-regulated genes in this cluster immediately underwent strong up-regulation and reached their highest expression one to two hours after peptidoglycan injection. By contrast, Toll-regulated genes underwent a delayed up-regulation of smaller magnitude, and reached their highest expression two to four hours after injection. Overall, the Bayesian LL*R*^2^ method successfully grouped together functionally related genes with distinct activation kinetics in this cluster.

In addition to recovering known dynamics of immune response pathways in cluster 10, the Bayesian LL*R*^2^ metric identified several new relationships between genes. In this cluster, four genes (*CG44404*, also known as *IBIN* ; *CG43236*, also known as *Mtk-like*; *CG43202*, also known as *BomT1* ; and *CG43920*) had no prior information available in the STRING database to link them with Imd-regulated genes. However, these four genes exhibit similar expression patterns to Imd-regulated genes, as seen in Figure 6, although previous studies examine *CG44404/IBIN* and *CG43202/BomT1* expression downstream of Toll signaling [Clemmons et al., 2015, Valanne et al., 2019]. This suggests that these four genes are not exclusively controlled by Toll signaling, and that they can also respond to Imd signaling after a gram-negative immune challenge. While Imd-regulation of *CG43236/Mtk-like* and *CG43920* has not been experimentally validated, their co-clustering pattern with Imd-regulated genes was also observed by Schlamp et al. [2021].

In Figure 7, we show a network consisting of the four aforementioned genes (*CG44404/IBIN, CG43236/Mtk-like, CG43202/BomT1, CG43920*) and their neighbors, i.e. the genes with which they have a Bayesian LL*R*^2^ of at least 0.9. Red edges in Figure 7 connect genes that were known to be associated according to prior information, i.e. **W**_*ij*_ = 1 for these pairs. Blue edges connect genes with an uncharacterized relationship in the STRING database, i.e **W**_*ij*_ = NA for these pairs.

**Figure 7:**
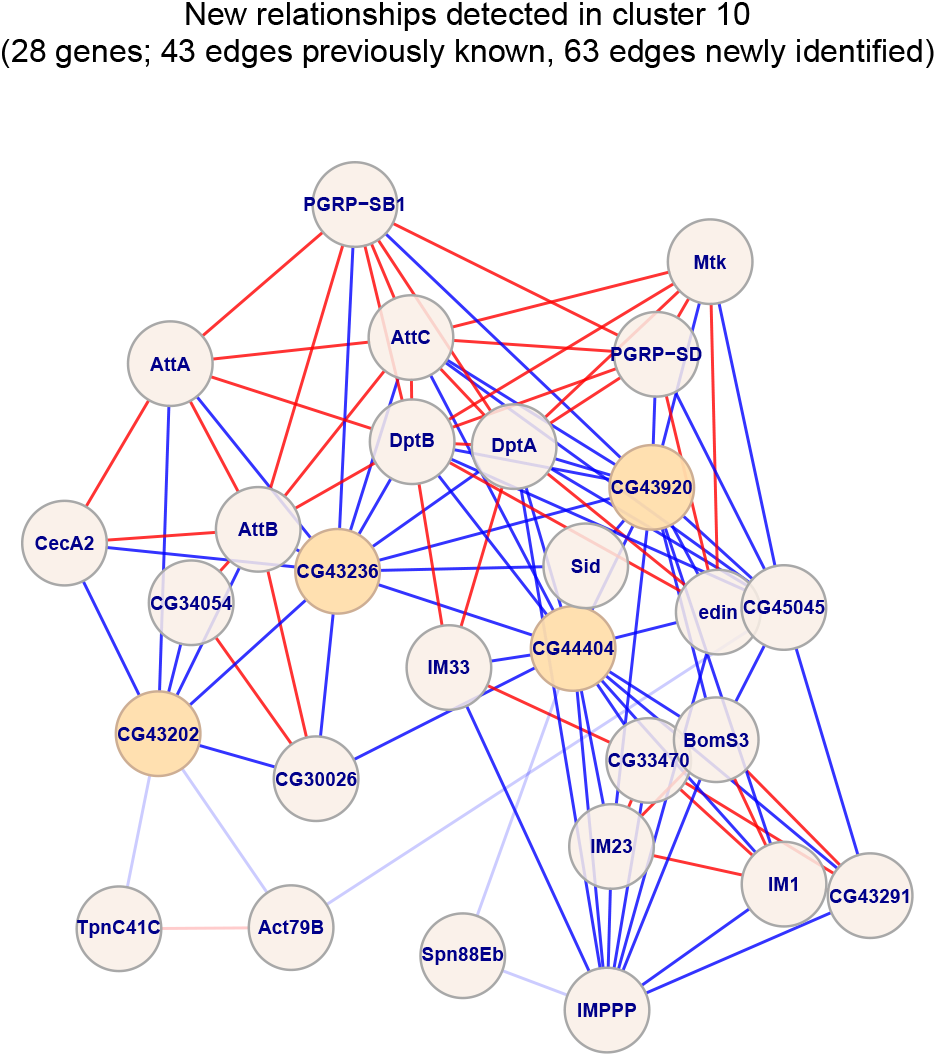
Network of genes formed from CG44404/IBIN, CG43236/Mtk-like, CG43202/BomT1, CG43920, and their neighbors. Two genes are connected by an edge if their Bayesian LLR^2^ > 0.9. Red and blue edges connect genes with known and unknown relationships, respectively. Darkened edges connect genes within cluster 10. Blue edges connect the four genes of interest with genes known to be regulated by the Imd signaling pathway, suggesting a possible role for them in fighting gram-negative infections.

### 4.4 The Bayesian LL*R*^2^ produces a sparse set of associations

The lead-lag *R*^2^ can be computed with biological information via our proposed Bayesian methodology, or without such information via ordinary least-squares regression. We now examine how the Bayesian approach changes the distribution of this quantity in a way that is conducive to identifying pairs or groups of genes with highly similar time dynamics.

In Section 3.5, we introduced the metrics 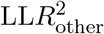 and 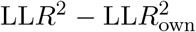. As described therein, the former indicates how much variation in gene *A*’s expression can be explained only through the dynamics of another gene *B*. The latter indicates how much additional variation in gene *A* can be explained by considering gene *B*, on top of considering gene *A*’s own past trajectory. Intuitively, both of these quantities should be large if the two genes are indeed associated in a way that manifests in highly similar temporal dynamics.

In Figure 8, we randomly select 150 genes and place all 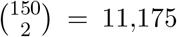 pairs on two scatterplots whose horizontal axes display the 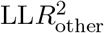 values and vertical axes display the 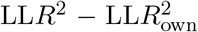 values. These *R*^2^ values are computed via our Bayesian method in one scatterplot and via ordinary least-squares regression in the other. Gene pairs of particular interest fall into the upper-right quadrant of the scatterplot.

**Figure 8:**
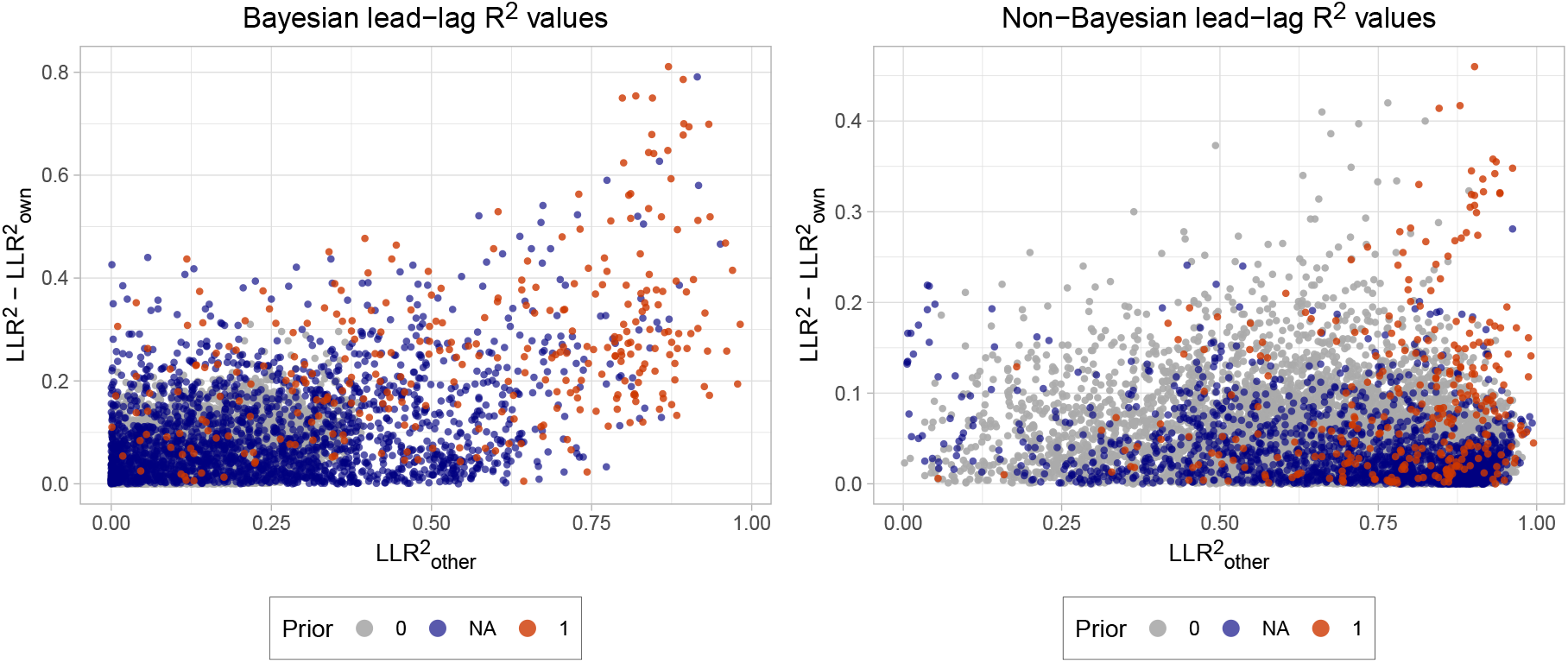
Scatterplots of 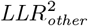 (horizontal axis) and 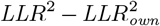 (vertical axis) for a random selection of 150 genes, resulting in 11,175 gene pairs. Left: LLR^2^ values are computed with the proposed Bayesian approach. Points are colored according to how prior information characterizes the corresponding two genes: unlikely to be associated (gray); uncharacterized association (blue); or known association (red). Right: LLR^2^ values are computed via ordinary least-squares, without external biological information.

Figure 8 shows that when we use the ordinary least-squares approach, i.e. without incorporating external biological information, we obtain an overwhelmingly large number of gene pairs with high 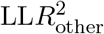 scores. Many are those that are unlikely to be associated, according to our chosen sources of prior information. By contrast, the Bayesian approach leverages this information to shift the distribution of the *R*^2^ values noticeably, yielding a smaller set of gene pairs that are worth examining further. This distributional shift is due to both the estimator ***β***_*_ in (10) for the coefficients in the model (4), as well as the way in which the parameter *g* is set in (15). In particular, *g* controls how much ***β***_*_ is influenced by either the data or prior information.

Importantly, Figure 8 shows that gene pairs with previously uncharacterized relationships but highly similar time dynamics are more easily identified with the Bayesian LL*R*^2^. A few of these gene pairs, which fall in the fairly sparse upper-right region of the left-hand scatterplot, are shown in Figure 9 in Appendix D. Figure 10 shows gene pairs in the same region of the scatterplot with well-known relationships, further demonstrating that the proposed method successfully recovers familiar associations.

**Figure 9:**
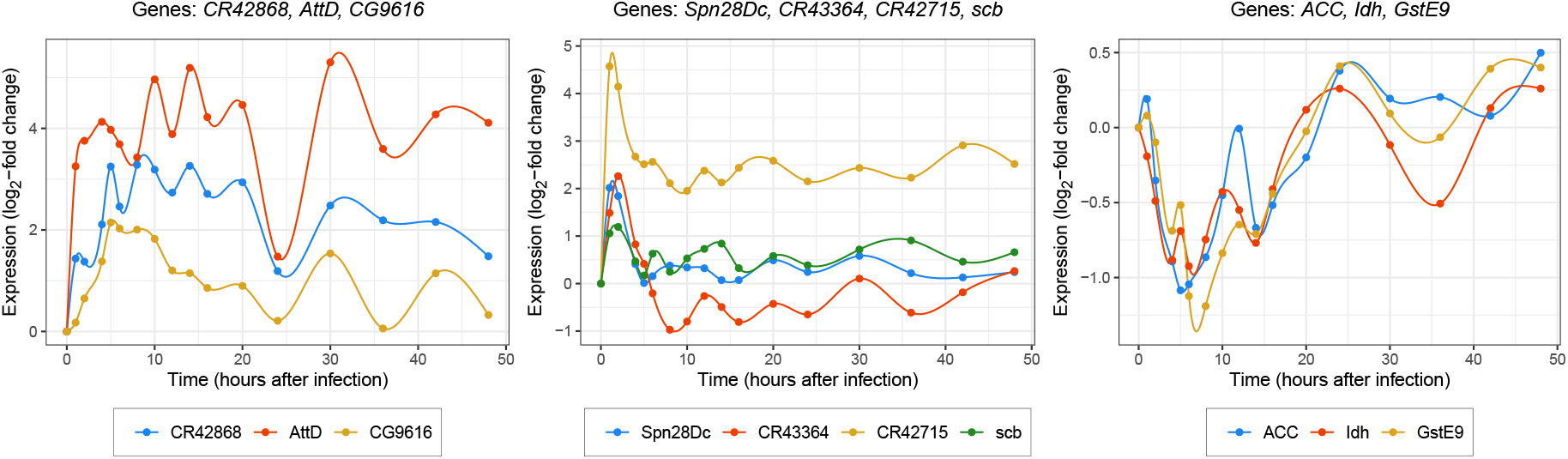
Sets of genes in the upper-right region of the Bayesian LLR^2^ scatterplot in Figure 8, several of which have uncharacterized pairwise associations. Left: AttD is involved in immune response against Gram-negative bacteria; it exhibits similar patterns to CR42868 and CG9616, neither of which have known molecular functions according to FlyBase [Larkin et al., 2020]. Middle: Spn28Dc is involved in response to stimuli and protein metabolism, and scb is involved in cell death and organ development. These two genes display similar expression patterns that are also nearly identical in shape to those of the less well-understood RNAs CR43364 and CR42715. Right: Genes ACC, Idh, and GstE9 are involved in a variety of metabolic processes, but not all of their pairwise interactions are known in STRING.

**Figure 10:**
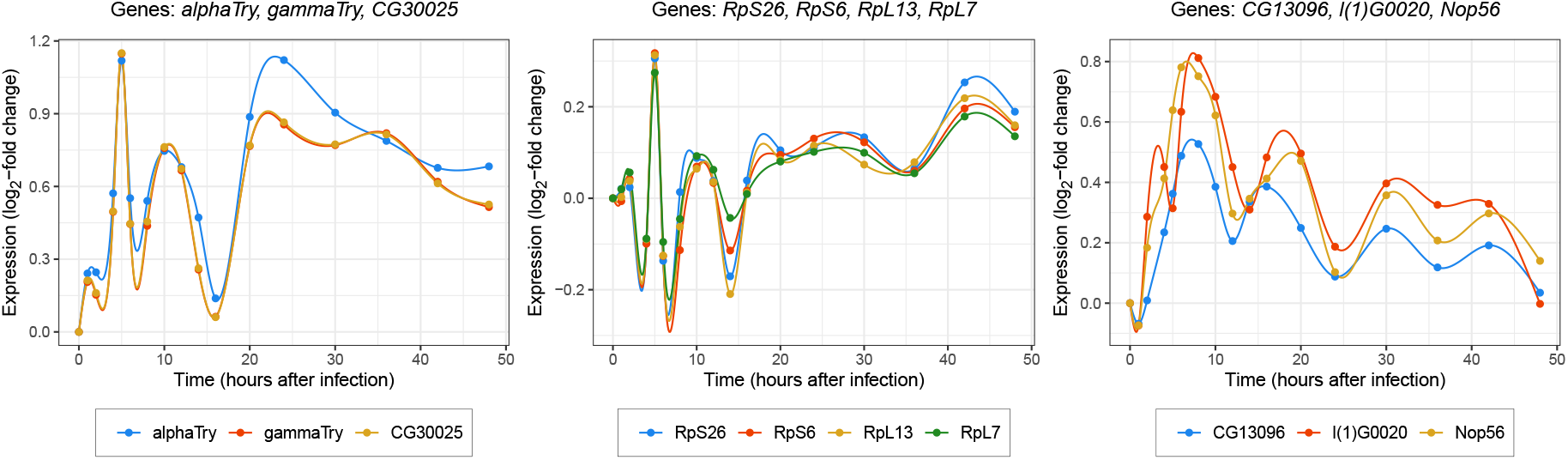
Sets of genes appearing in the upper-right region of the Bayesian LLR^2^ scatterplot in Figure 8, all of which have known pairwise associations. Left: Genes known to be involved in proteolysis. Middle: Genes that encode ribosomal proteins. Right: Genes known to be involved in RNA binding.

**Figure 11:**
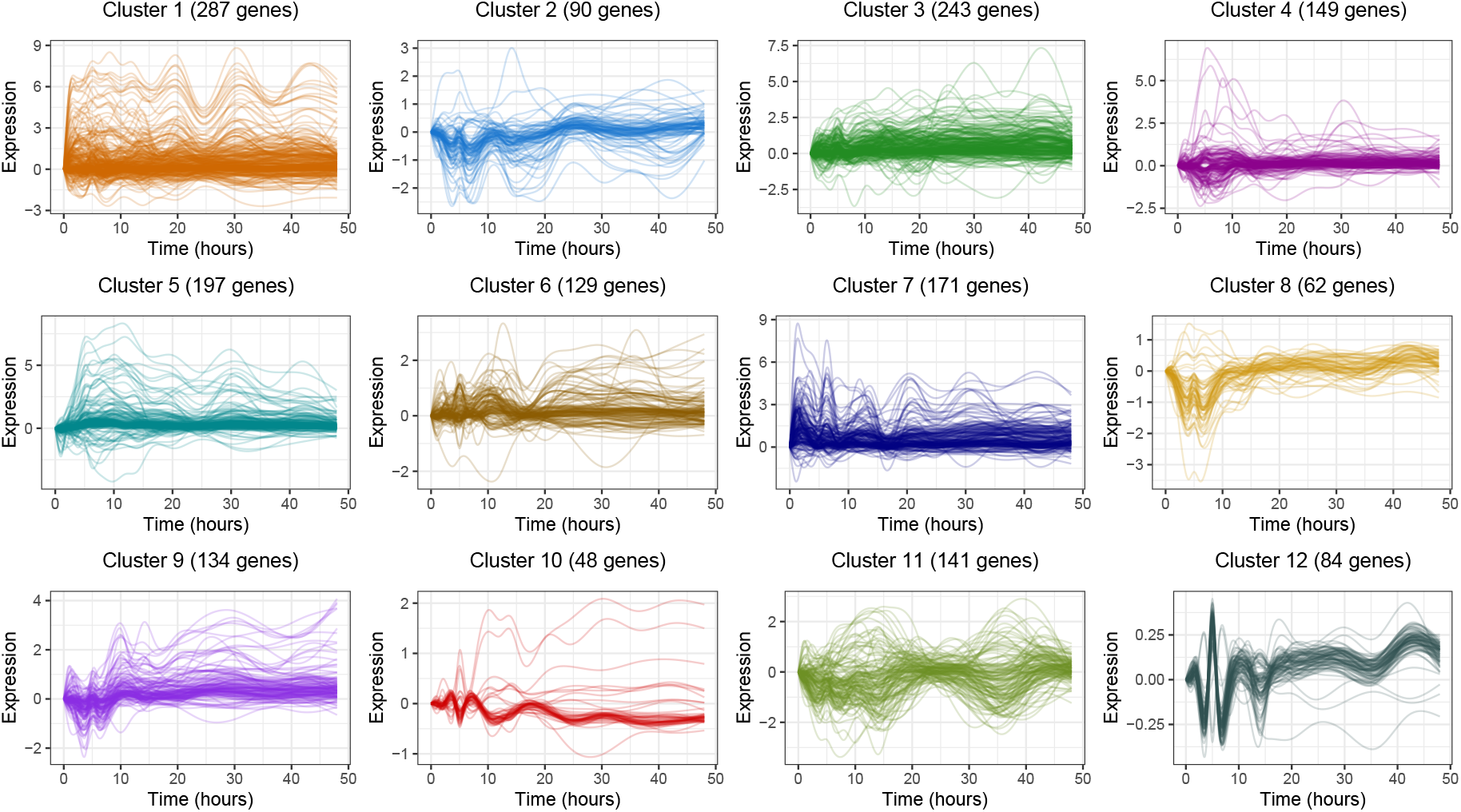
Clusters of genes obtained using the non-Bayesian LLR^2^ as the similarity metric between genes. Each sub-plot shows the temporal expressions of genes in the corresponding cluster. Clusters were obtained via hierarchical clustering with Ward’s method. The dominant pattern in many clusters is either flat or not immediately discernible, indicating that the non-Bayesian LLR^2^ is alone insufficient for identifying groups of genes whose temporal patterns are of similar shapes over time.

**Figure 12:**
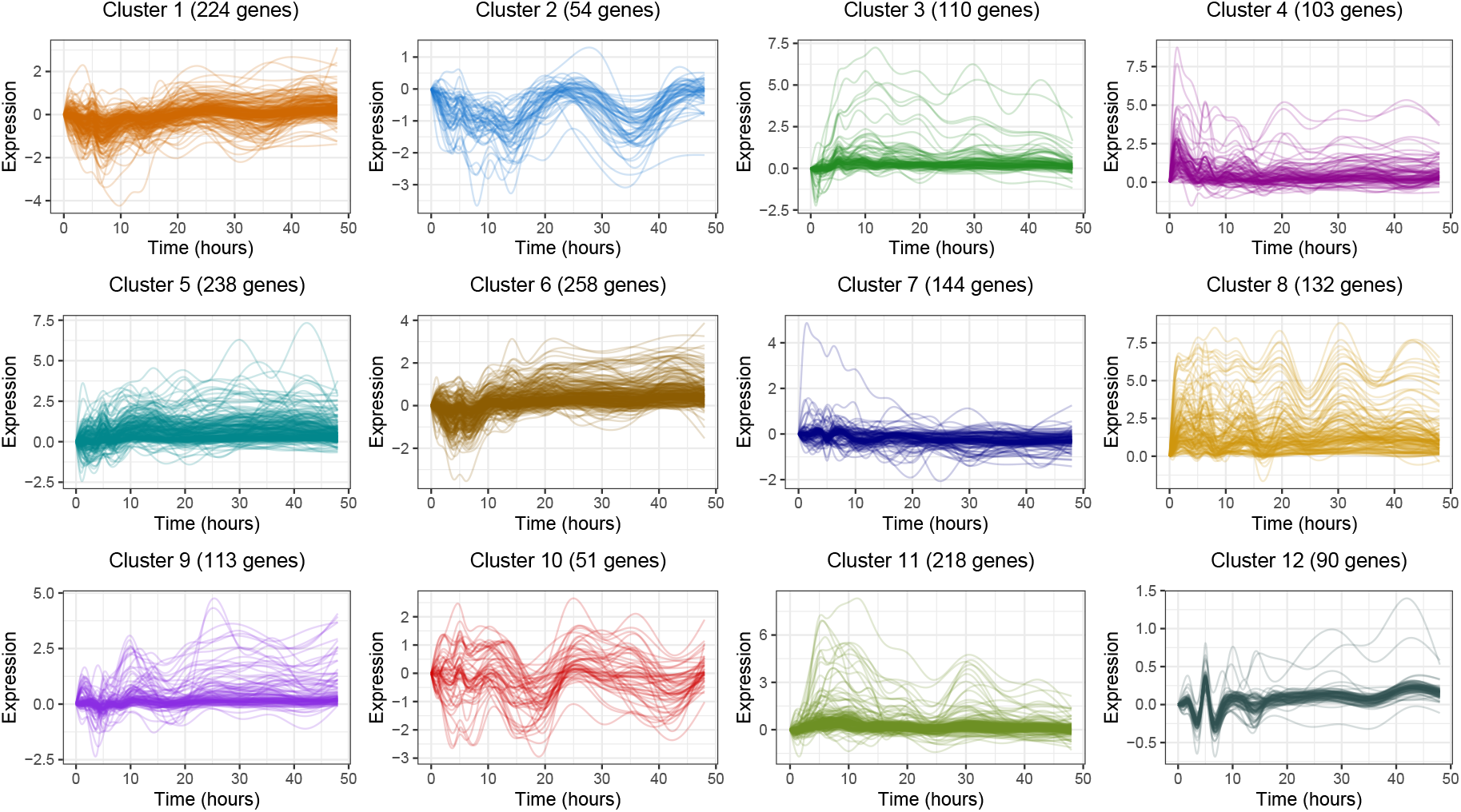
Clusters of genes obtained using Pearson correlation as the similarity metric between genes. Each sub-plot shows the temporal expressions of genes in the corresponding cluster. In cluster and network analyses of time-course gene expression data, Pearson correlation is one of the more commonly used metrics of association. Clusters were obtained via hierarchical clustering with Ward’s method. The dominant pattern in many clusters is either flat or not immediately discernible, indicating that correlation is alone insufficient for identifying groups of genes whose temporal patterns are of similar shapes over time.

## 5 Discussion

Time-course gene expression datasets are a valuable resource for understanding the complex dynamics of interconnected biological systems. An important statistical task in the analysis of such data is to identify clusters or networks of genes that exhibit similar temporal expression patterns. These patterns yield systems-level insight into the roles of biological pathways and processes such as disease progression and recovery.

The main statistical challenges in studying time-course gene expression datasets stem from their high dimensionality and small sample sizes, combined with the nonlinearity of gene expression time dynamics. To overcome these difficulties, we proposed a method for identifying potentially associated genes that treats temporal gene expression as a dynamical system governed by ODEs, whose parameters are determined in a Bayesian way using gene networks curated *a priori* from biological databases. The ODE model is fit to a pair of genes via Bayesian regression and is used to derive the biologically-informed Bayesian lead-lag *R*^2^ similarity measure. The Bayesian regression procedure leverages Zellner’s *g*-prior and ideas from shrinkage estimation, namely minimization of Stein’s unbiased risk estimate (SURE), to balance the posterior ODE model’s fit to the data and to prior information. As a result, we automatically encourage gene pairs with known associations to receive higher Bayesian lead-lag *R*^2^ scores, while reducing the scores of gene pairs that are unlikely to be related and allowing new relationships to be discovered.

In Section 4, we analyzed clusters and networks of genes that were identified by our method as having similar temporal dynamics. In particular, the clusters highlighted the known interplay between immune response and metabolism, and suggested roles for uncharacterized genes displaying remarkably similar temporal patterns to more well-studied ones. We contrasted our results to those obtained by using only the ordinary least-squares version of the lead-lag *R*^2^ and demonstrated how the inclusion of prior biological information greatly aids the identification of biologically relevant gene groups.

## A Proofs

In this section, we provide proofs of Theorems 3.2 and 3.3.

*Proof of Theorem 3.2*. We write *δ*_0_(**Y**) as *δ*_0_(*g*), treating **Y** as fixed. Expanding the expression for *δ*_0_(**g**), we obtain

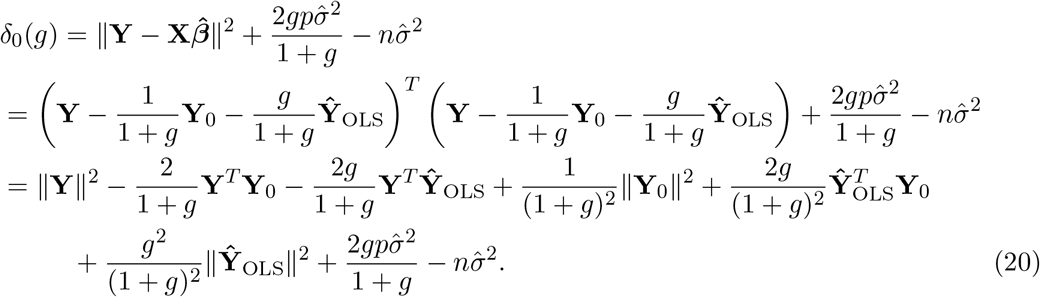

Next, observe that **Y**^*T*^**Ŷ**_OLS_ = ‖**Ŷ**_OLS_‖^2^, because

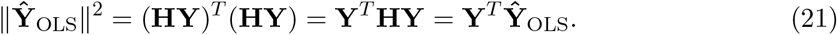

Furthermore, observe that 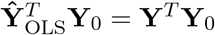, because

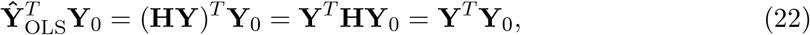

where the last equality follows from the fact that **Y**_0_ = **X*β***_0_, i.e. **Y**_0_ is in the column span of **X**, which is the space onto which **H** projects. The identities (21) and (22) can now be used to write *δ*_0_(*g*) in (20) as

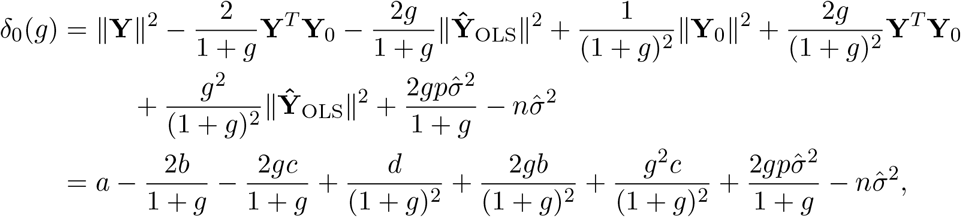

where *a* = ‖**Y**‖^2^, *b* = **Y**^*T*^**Y**_0_, *c* = ‖**Ŷ**_OLS_‖^2^, and *d* = ‖**Y**_0_‖^2^. Differentiating *δ*_0_(*g*) with respect to *g*, we obtain

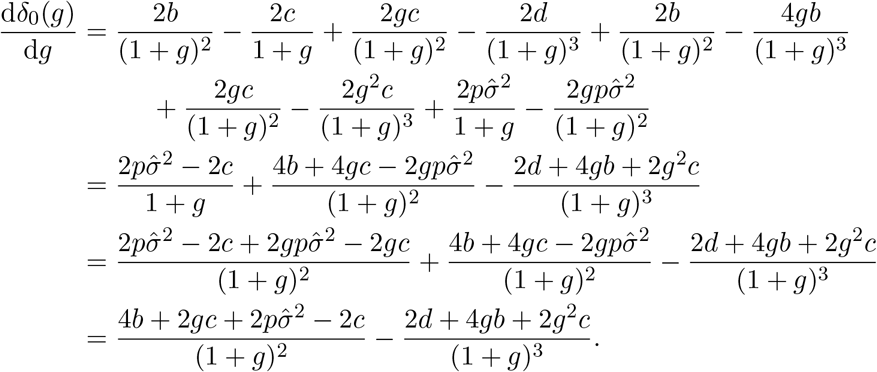

Setting this derivative to zero and rearranging yields:

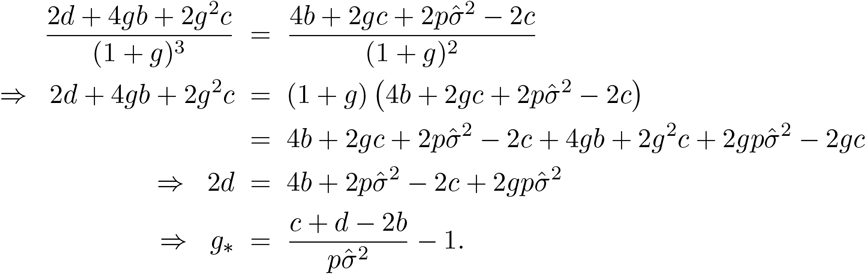

We now substitute the definitions of *b, c*, and *d* back into this expression to obtain

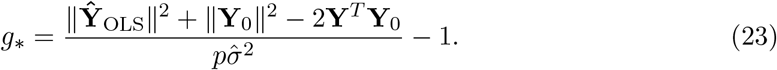

The numerator of (23) can be simplified by observing that

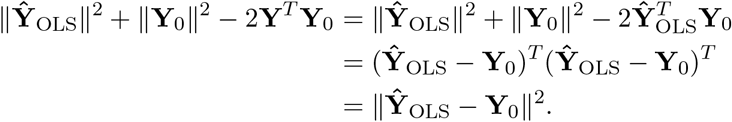

Therefore, (23) becomes

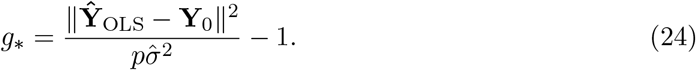

When ***β***_0_ = **0**, we have **Y**_0_ = **X*β***_0_ = **0**. In this case, (24) becomes

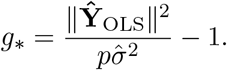

Finally, the second derivative of *δ*_0_(*g*) evaluated at *g* = *g*_*_ in (24) is equal to

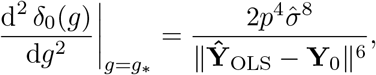

which is positive, thus confirming that *δ*_0_(*g*) is indeed minimized at *g* = *g*_*_. This second derivative calculation was verified in Mathematica.□

*Proof of Theorem 3.3*. We write *δ*_0_(**Y**) as *δ*_0_(*g, ξ*), treating **Y** as fixed. First, if ***β***_0_ = [*ξ, ξ*, 0, 0, 0]^*T*^, then **X*β***_0_ = *ξ***X**_12_, where **X**_12_ is the element-wise sum of the first two columns of **X**. We now proceed similarly to the proof of Theorem 3.2 by expanding *δ*_0_(*g, ξ*):

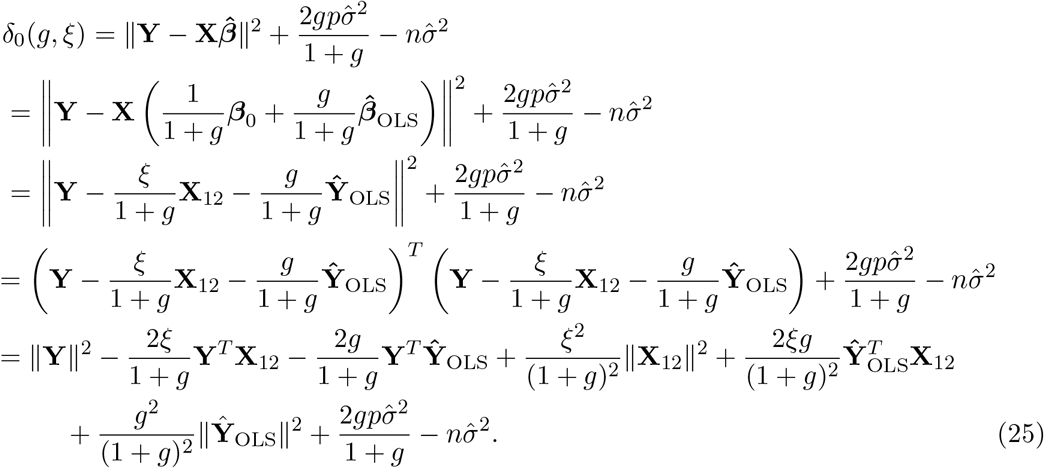

Next, observe that 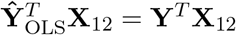, because

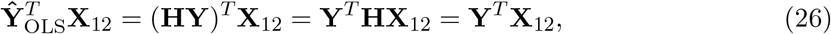

where the last equality follows from the fact that **X**_12_ is in the column span of **X**, which is the space onto which **H** projects. The identities (21) and (26) can now be used to write *δ*_0_(*g, ξ*) in (25) as

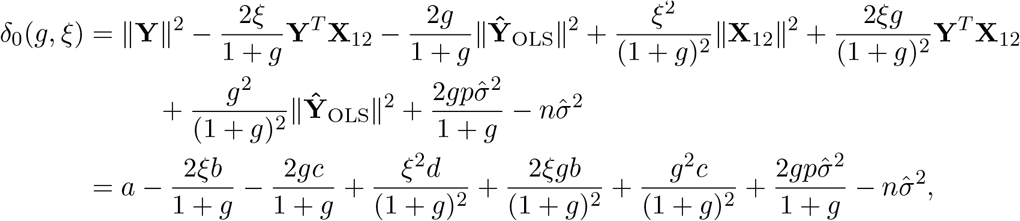

where *a* = ‖**Y**‖^2^, *b* = **Y**^*T*^**X**_12_, *c* = ‖**Ŷ**_OLS_‖^2^, and *d* =‖ **X**_12_‖^2^. Differentiating *δ*_0_(*g, ξ*) with respect to *ξ*, we obtain

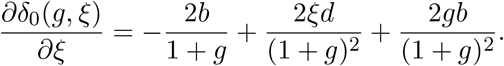

Setting this derivative to zero and rearranging yields:

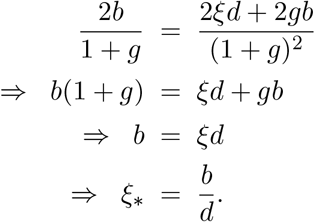

We now substitute the definitions of *b* and *d* back into this expression to obtain

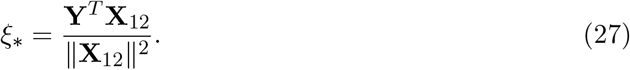

Next, we differentiate *δ*_0_(*g, ξ*) with respect to *g*:

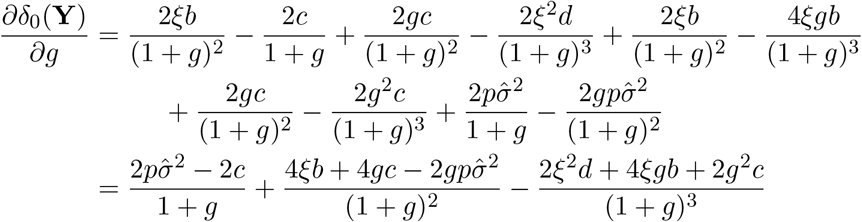

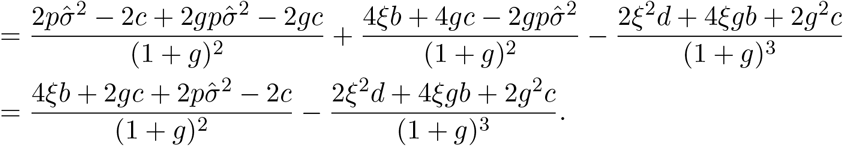

Setting this derivative to zero and rearranging yields:

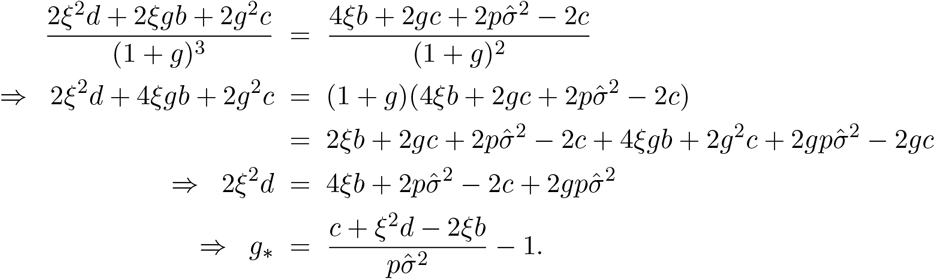

We now substitute the definitions of *b, c*, and *d* back into this expression to obtain

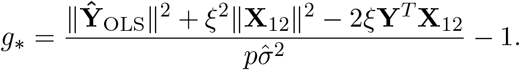

Substituting *ξ* in (27) into this expression for *g* yields

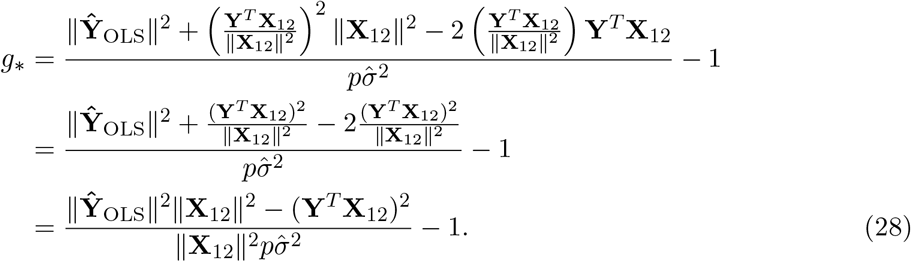

Finally, the determinant of the Hessian matrix of *δ*_0_(*g, ξ*), denoted ∇^2^*δ*_0_, evaluated at *g* = *g*_*_ in (28) and *ξ* = *ξ*_*_ in (27) is

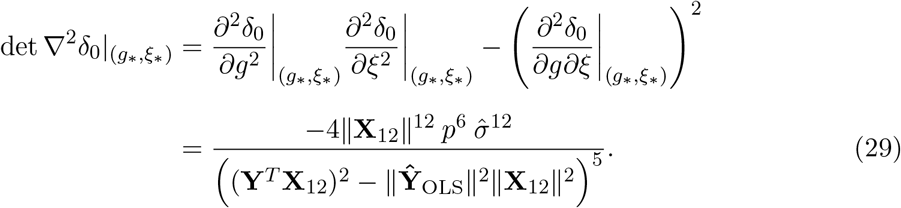

We must now verify that det 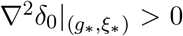, i.e. that (*g*_*_, *ξ*_*_) is indeed an extremum of *δ*_0_. First, we observe that

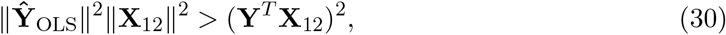

by direct application of the Cauchy-Schwarz inequality (recall that 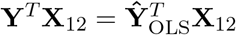 from (26)). The inequality is strict because of the assumption that the entries of 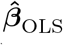 are distinct, which prevents **Ŷ**_OLS_ and **X**_12_ from being linearly dependent. Thus, (30) implies that the denominator in (29) is strictly negative. The numerator is also strictly negative, because the assumption that gene *B* is not zero everywhere ensures that ‖**X**_12_‖ ^2^ ≠ 0 (recall that **X** is defined in (5), and its first two columns consist of gene *B*’s expression measurements over time and its time integrals). Therefore, (29) is strictly positive. To verify that (*g*_*_, *ξ*_*_) is indeed a minimizer of *δ*_0_, we now check that 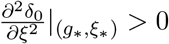 as well. We have:

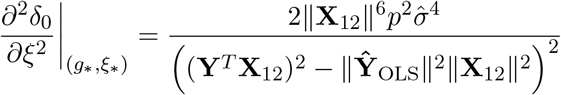

which is strictly positive. This second derivative calculation, as well as those in (29), were verified in Mathematica.□

## B Bayesian lead-lag *R*^2^ algorithm

Following is an algorithm for computing the Bayesian lead-lag *R*^2^ for all gene pairs in a time-course gene expression dataset, consisting of *N* genes measured at *n* time points *t*_1_, …, *t*_*n*_.

### Algorithm 1: Bayesian lead-lag *R*^2^ calculations for all gene pairs

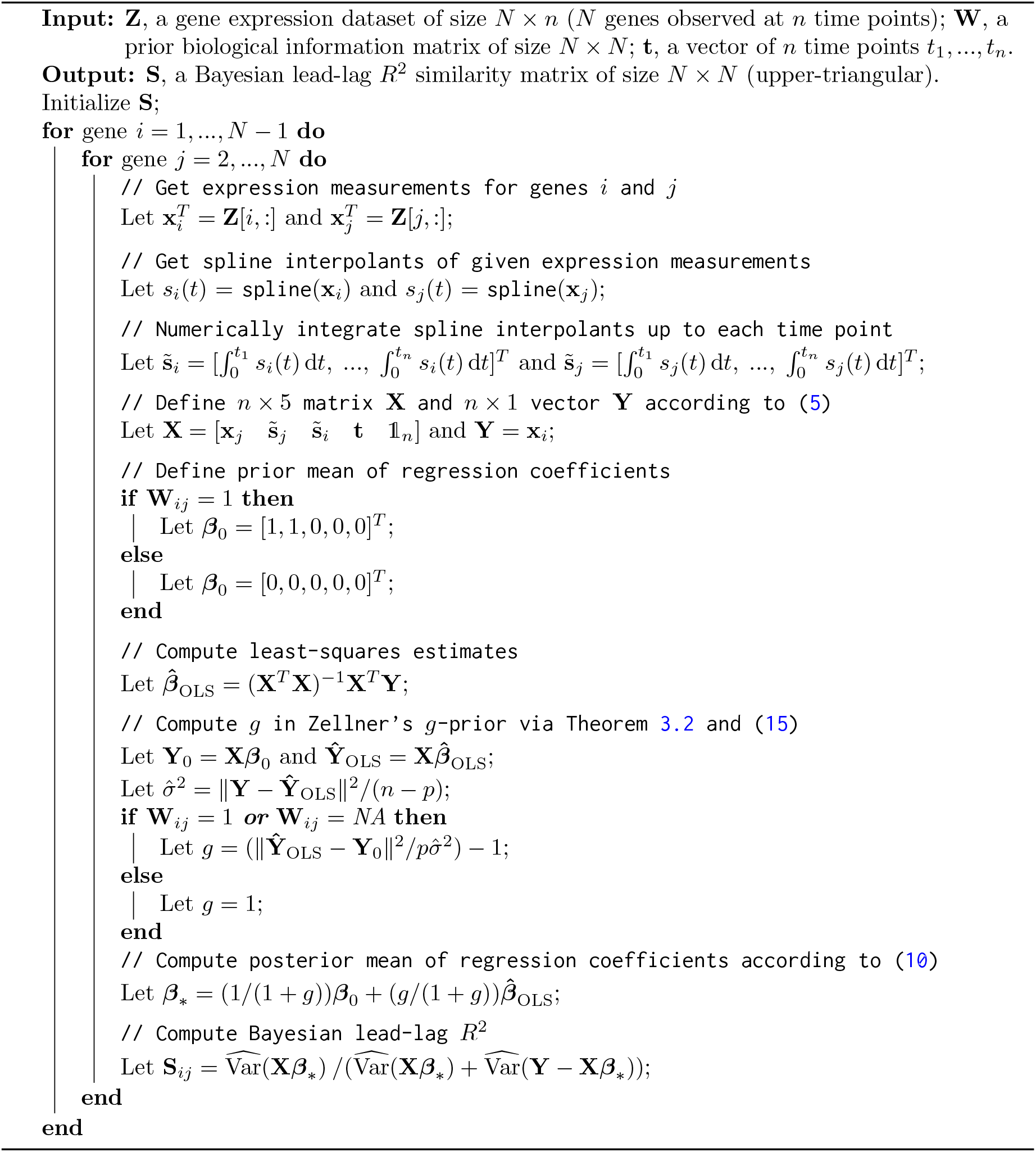

Note that this algorithm produces an upper-triangular similarity matrix **S** resulting from regressing gene *i* on gene *j*, for all *i < j*, and storing the resulting Bayesian LL*R*^2^(*i, j*) value. In the empirical analysis presented in this paper, we set **S**_*i,j*_ = max{LL*R*^2^(*i, j*), LL*R*^2^(*j, i*)}.

To instead compute the Bayesian 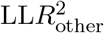 from sub-model 1 in (18), it suffices to change the definition of **X** to 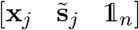, and to define ***β***_0_ as [1, 1, 0]^*T*^ if **W**_*ij*_ = 1 and [0, 0, 0]^*T*^ otherwise. To compute the Bayesian 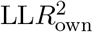 from sub-model 2 in (19), we change **X** to 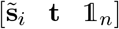, and ***β***_0_ = [0, 0, 0] regardless of the value of **W**_*ij*_. An optimized version of this algorithm runs in 21.8 minutes for *N* = 1735 genes on a 2017 3.1 GHz Intel Core i5 MacBook Pro.

## C Additional dataset details

In this section, we provide further details on how our primary time-course gene expression dataset was constructed.

We first reduce our set of 12,657 genes into a set of 951 “differentially-expressed” (DE) genes identified in Schlamp et al. [2021], defined as genes satisfying either of the following criteria: 1) there is at least one time point at which the gene’s expression undergoes a log_2_-fold change of at least two, or 2) a spline-based method for differential expression analysis returns a significant result. The latter method involves fitting a cubic spline to the temporal expression measurements of a gene under treatment and control settings, and testing whether the difference in the resulting two sets of coefficients is significant. We then add back a set of 784 genes that are not DE by these criteria, but that are “neighbors” of at least one DE gene. We define a neighbor of a DE gene as a non-DE gene that has a STRING score of at least 0.95 with the DE gene. The purpose of adding such neighbors back into the dataset, now consisting of 951 + 784 = 1735 genes, is to enable more complete biological pathways to be reconstructed from our cluster analysis.

In Section 3.1, we describe several sources of biological information that can be encoded into a prior adjacency matrix **W**. We choose to use the STRING database, and we mark two genes as “associated” (i.e., **W**_*ij*_ = 1) if their STRING score is greater than 0.5. We additionally use replicate information from the time-course dataset to fill in some of the unknown STRING scores. Specifically, if two genes have entries in the STRING database but have an unknown STRING score, we set **W**_*ij*_ = 1 if the correlation between their replicated temporal expressions is greater than 0.8. We keep **W**_*ij*_ = NA for the gene pairs that do not have entries in STRING.

## D Additional figures

## E Additional tables

## F Additional results

We continue our analysis in Section 4.3 of the results of hierarchical clustering with the Bayesian lead-lag *R*^2^ (LL*R*^2^).

### F.1 Analysis of cluster 2

Cluster 2 contains both up- and down-regulated genes with circadian rhythms, according to GO term enrichment. Several of these genes are displayed in Figure 13. Among the up-regulated genes are three regulators of the circadian clock (*per, vri, Pdp1*). A fourth regulator of the circadian clock, *Clk*, is down-regulated. *Pdp1* has been reported to reach its peak expression three to six hours after *vri* ‘s peak expression [Cyran et al., 2003], a pattern that is visible in this cluster. Cluster 2 further contains genes that are involved in visual perception: two genes encoding rhodopsins (*Rh5, Rh6*) [Gaudet et al., 2011] and *Pdh*, which encodes a retinal pigment dehydrogenase [Wang et al., 2010]. Similar to Schlamp et al. [2021], we also found that *Smvt* and *salt*, which encode sodium transporters [Gaudet et al., 2011], are under circadian control.

**Figure 13:**
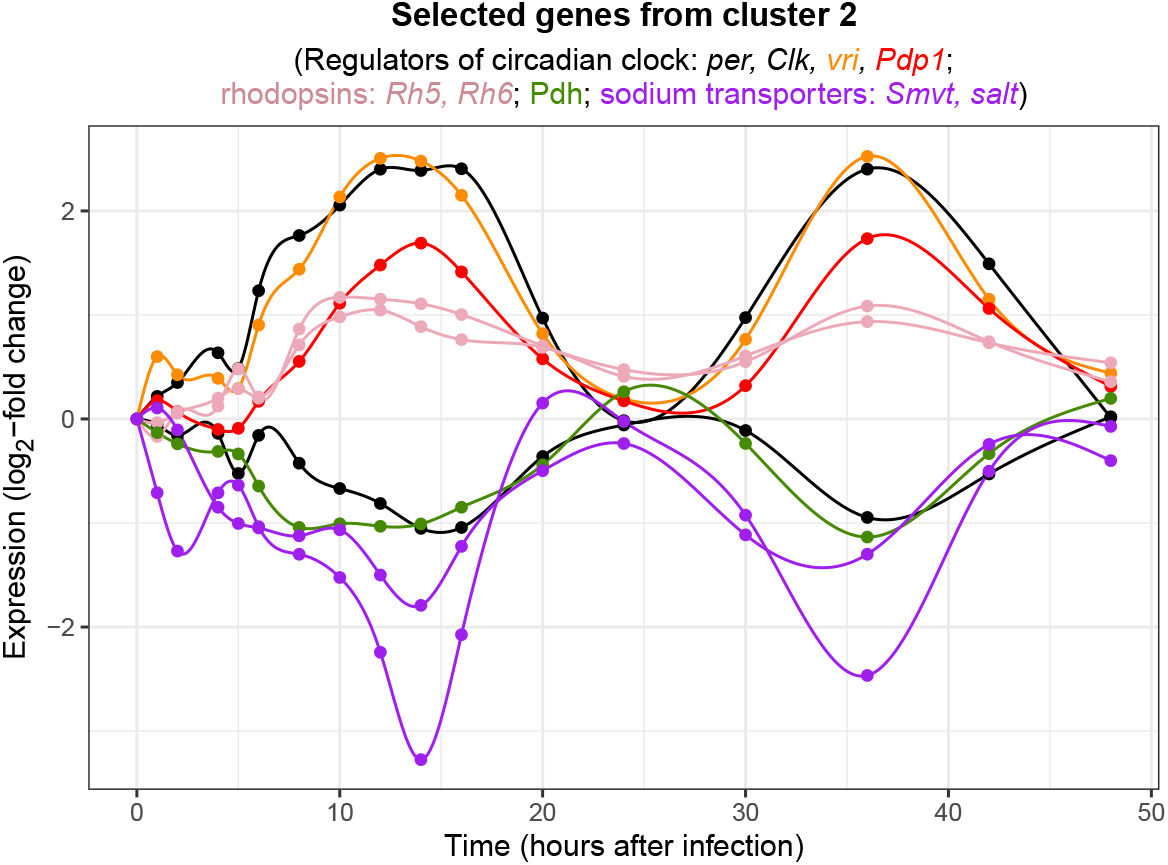
Temporal expression patterns of selected genes in cluster 2. Black, red, and orange lines correspond to regulators of the circadian clock (black: per, Clk; orange: vri; red: Pdp1). Pdp1 is known to reach its peak expression after vri does. Pink and green lines correspond to genes that are involved in visual perception (pink: rhodopsins Rh5, Rh6; green: retinal pigment dehydrogenase Pdh). Purple lines correspond to genes that encode sodium transporters (Smvt, salt).

### F.2 Analysis of cluster 4

Genes in cluster 4, some of which are displayed in Figure 14, are characterized by a transient decrease in expression during the first 24 hours after peptidoglycan injection. This cluster was significantly enriched for “carbohydrate metabolic process” (B-H corrected *p*-value of 2 × 10^−23^). A highly-connected gene involved in carbohydrate metabolism is *fbp*, which encodes the enzyme fructose-1,6-biphosphatase and has a degree of 102 in our reconstructed network shown in Figure 15.

**Figure 14:**
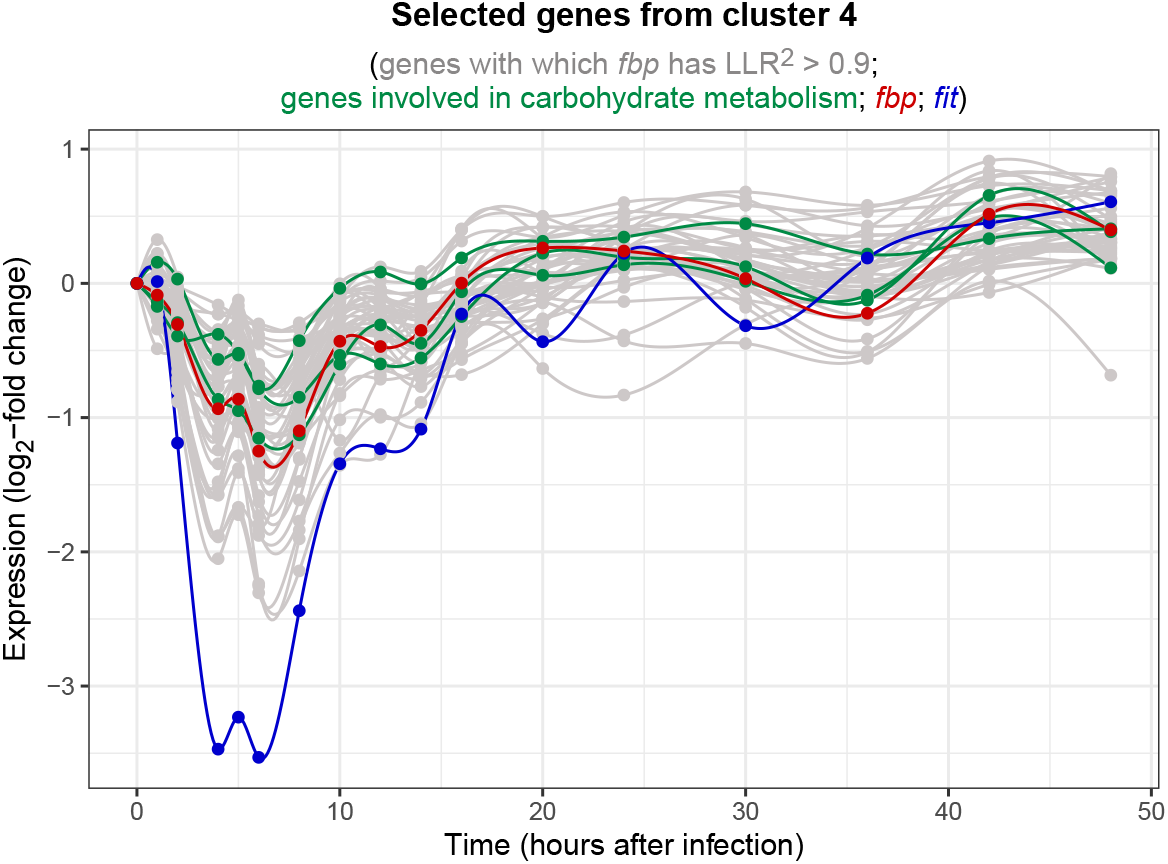
Temporal expression patterns of selected genes in cluster 4. Light gray lines in the background correspond to genes with which the gene fbp has a Bayesian LLR^2^ > 0.9. fbp, shown in red, is known to be involved in carbohydrate metabolism. The three green lines correspond to genes Gale, AGBE, and Gba1b, which also have known roles in carbohydrate metabolism but have an unknown relationship to fbp according to the STRING database. The dark blue line corresponds to the gene fit, whose expression pattern is similar to that of fbp but with much more pronounced down-regulation. fit is known to encode a protein that stimulates insulin signaling, a process that regulates the expression of genes involved in carbohydrate metabolism.

**Figure 15:**
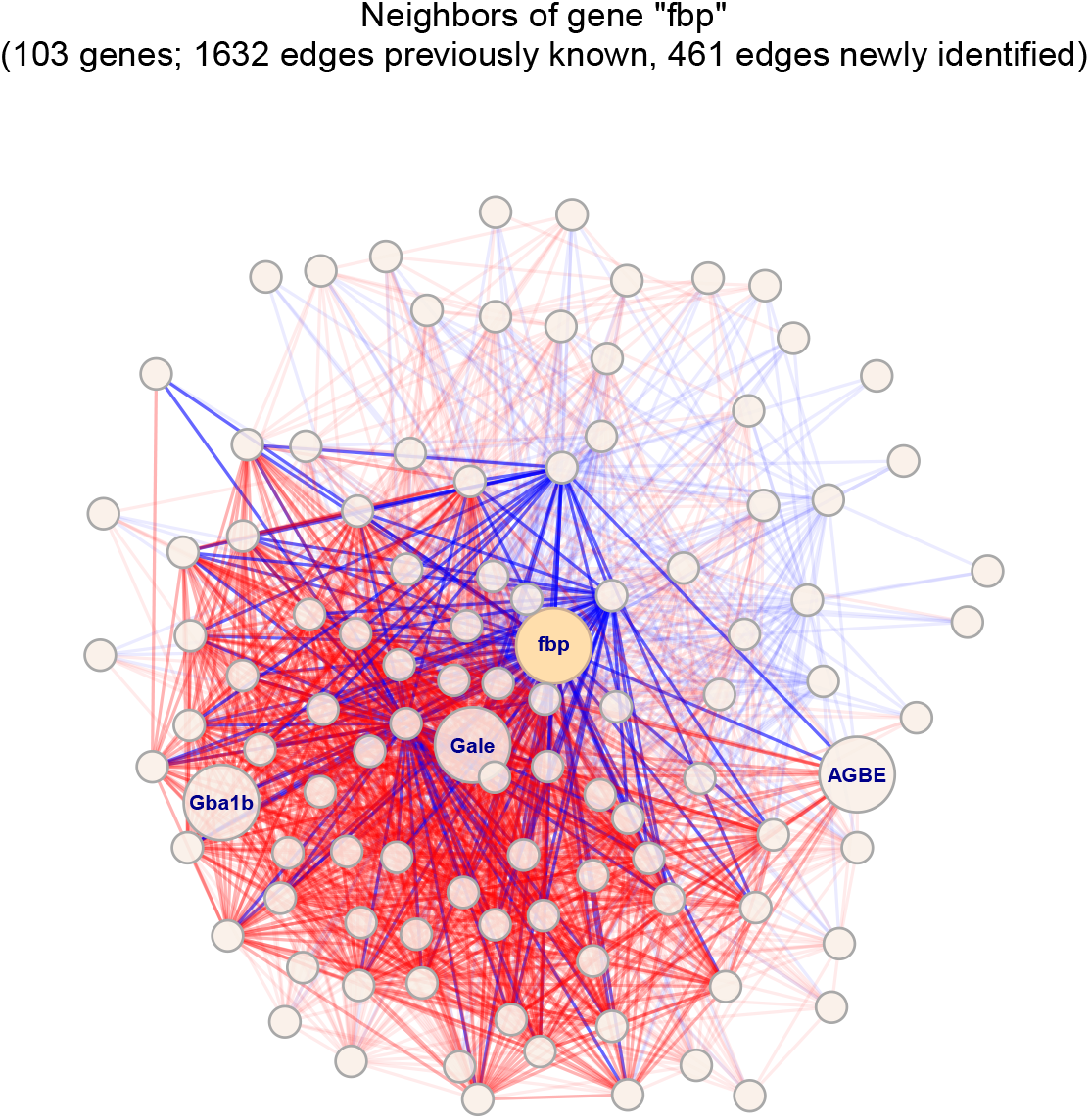
Network of genes formed from the gene fbp and its neighbors. Two genes are connected by an edge if their Bayesian LLR^2^ > 0.9. Red and blue edges connect genes with known and unknown relationships, respectively. Darkened edges connect genes within cluster 4. Selected genes with uncharacterized relationships to fbp are highlighted: Gale, AGBE, Gba1b.

All of the genes to which *fbp* is connected had NA values in our prior adjacency matrix, meaning that their relationship to *fbp* is unknown according to our chosen sources of information. Some of these genes include *Gale, AGBE*, and *Gba1b*, which have known roles in carbohydrate metabolism. Twenty-two other genes connected to *fbp* in our network are not well-studied. These connections suggest roles for these uncharacterized genes in carbohydrate metabolism or energy homeostasis. Another gene connected to *fbp* is *fit*, which has a similar expression profile as *fbp* but experiences a much stronger and sharper down- and up-regulation. *fit* is not directly involved in carbohydrate processing but encodes a secreted protein that stimulates insulin signaling, which in turn regulates the expression of genes involved in carbohydrate metabolism, such as *fbp* [Sun et al., 2017]. A previous study also showed that an immune response reduces insulin signaling in *Drosphila* [DiAngelo et al., 2009].

### F.3 Analysis of cluster 6

Cluster 6 is significantly enriched for GO terms related to metabolic processes, particularly the terms “cellular lipid catabolic process” and “carbohydrate metabolic process” (B-H corrected *p*-values of 2^−10^ for both). In addition to these metabolic GO terms, there is significant enrichment of genes involved in “phagocytosis” (B-H corrected *p*-value of 0.03). During an immune response, phagocytosis is the process by which an immune cell engulfs and digests bacteria and apoptotic cells as a way to fight the infection. In *Drosophila*, phagocytosis is carried out by specialized hemocytes (*Drosophila* blood cells).

Among the genes in cluster 6 involved in carbohydrate metabolism are five mannosidases (*LManI, LManIII, LManIV, LManV, LManVI*) and three maltases (*Mal-A2, Mal-A3, Mal-A4*). These genes encode enzymes that break down complex sugars into simple sugars like glucose. Six genes in cluster 6 are expressed in hemocytes and are involved in phagocytosis. These include four genes that belong to the Nimrod gene family (*NimB4, NimC1, NimC2, eater*); *Hml*, which is involved in the clotting reaction in larvae [Goto et al., 2003]; and *Gs1. Gs1* encodes a glutamine synthetase, an enzyme whose action is not unique to hemocytes, but that has been shown to support hemocyte function [Gonzalez et al., 2013].

Figure 16 shows that the genes involved in metabolic processes and in phagocytosis exhibit similar expression patterns, with coordinated up- and down-regulation. This coordinated expression is sensible in the context of known hemocyte biology. After an infection, hemocytes undergo a metabolic switch, whereby their energy production is sustained mostly by aerobic glycolysis rather than oxidative phosphorylation [Krejčová et al., 2019]. Since aerobic glycolysis is dependent on glucose, the simultaneous up-regulation of glucose-producing enzymes and genes needed for phagocytosis is aligned with our expectations. It is also worth noting that the fold changes of genes involved in phagocytosis are generally small, e.g. less than two-fold up-regulation, which is often used as a minimal cutoff in RNA-seq analyses. However, the coordinated expression changes detected by the Bayesian LL*R*^2^ suggest that these are biologically relevant patterns.

**Figure 16:**
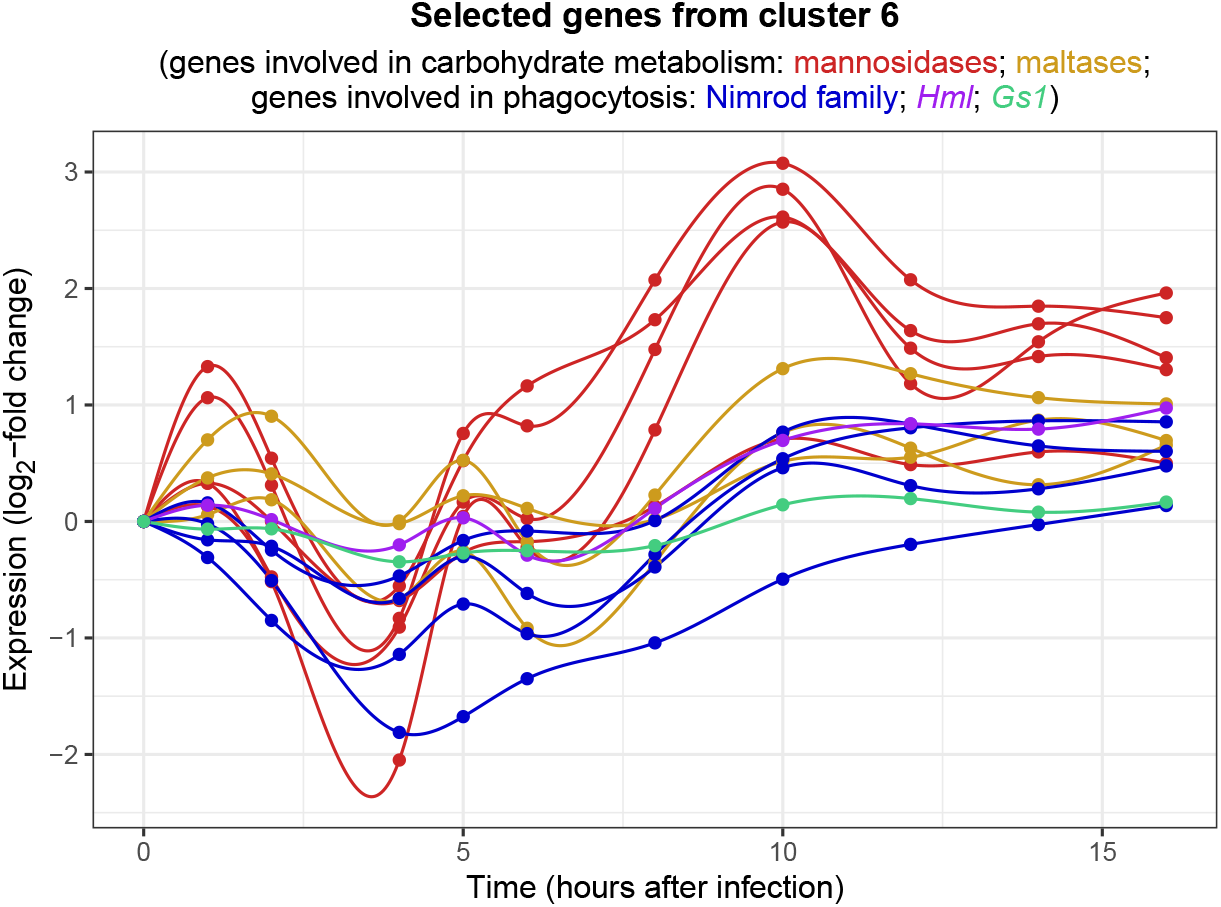
Temporal expression patterns of selected genes in cluster 6 during the first 16 hours after peptidoglycan injection. The five red lines and three yellow lines correspond to genes involved in carbohydrate metabolism: respectively, mannosidases (LMan1, LManIII, LManIV, LManV, LManVI) and maltases (Mal-A2, Mal-A3, Mal-A4). The remaining lines correspond to genes that are expressed in hemocytes and are involved in phagocytosis: in blue are genes belonging to the Nimrod family (NimB4, NimC1, NimC3, eater), in purple is Hml, and in green is Gs1.

